# Self-extinguishing relay waves enable homeostatic control of human neutrophil swarming

**DOI:** 10.1101/2023.06.27.546744

**Authors:** Evelyn Strickland, Deng Pan, Christian Godfrey, Julia S. Kim, Alex Hopke, Maureen Degrange, Bryant Villavicencio, Michael K. Mansour, Christa S. Zerbe, Daniel Irimia, Ariel Amir, Orion D. Weiner

## Abstract

Neutrophils exhibit self-amplified swarming to sites of injury and infection. How swarming is controlled to ensure the proper level of neutrophil recruitment is unknown. Using an *ex vivo* model of infection, we find that human neutrophils use active relay to generate multiple pulsatile waves of swarming signals. Unlike classic active relay systems such as action potentials, neutrophil swarming relay waves are self-extinguishing, limiting the spatial range of cell recruitment. We identify an NADPH-oxidase-based negative feedback loop that is needed for this self-extinguishing behavior. Through this circuit, neutrophils adjust the number and size of swarming waves for homeostatic levels of cell recruitment over a wide range of initial cell densities. We link a broken homeostat to neutrophil over-recruitment in the context of human chronic granulomatous disease.

---

During collective migration, individual organisms coordinate their movement to solve critical tasks. Birds flock, fish school, and insects swarm to escape predation, find food, and move more efficiently Chandra et al. (2021); Gordon (2019); Parrish and Edelstein-Keshet (1999). These organismal collectives rely on individuals acting on local information, including distance to neighbors, alignment with neighbors, and local chemical cues, to generate complex emergent decisions. Collective migration is also organized at the cellular level, where groups of cells coordinate wound healing LaChance et al. (2022); Sun et al. (2023), cancer metastasis Muinonen-Martin et al. (2014), multicellular morphogenesis Donà et al. (2013), and the transition from single cell to multicellular existence Gregor et al. (2010). We are just beginning to understand the molecular logic of cell-cell communication that enables collective behaviors to arise. Here we probe the logic of collective migration in the context of the human immune response. Neutrophils are first responders of the innate immune system that are recruited to sites of injury and infection to neutralize invading pathogens and aid in tissue repair Ley et al. (2018); Kolaczkowska and Kubes (2013); Peiseler and Kubes (2019). Because the primary signals of injury/infection are relatively short ranged, activated neutrophils release chemoattractants such as Leukotriene B4 (LTB4) for positive feedback-based recruitment of additional neutrophils. This self-amplified ‘swarming’ process significantly enhances the speed and range of recruitment Kienle et al. (2021); Sun and Shi (2016); Poplimont et al. (2020); Reátegui et al. (2017); Lämmermann et al. (2013), but this process must be tightly regulated to limit collateral damage Kienle et al. (2021); Uderhardt et al. (2019); Jiwa et al. (2020); Mawhin et al. (2018). While some brakes on swarming are known Kienle et al. (2021); Reátegui et al. (2017); Uderhardt et al. (2019), how neutrophils control the spatiotemporal dynamics of cell-cell communication to recruit the appropriate number of cells is not well understood, and several fundamental questions remain unanswered. Are swarms primarily constrained by the neutrophils themselves Kienle et al. (2021), or are other cells required Uderhardt et al. (2019)? Do different molecular programs control the duration versus the range of swarming? And how robust is swarming to initial conditions?

While many neutrophil swarming studies have been performed in living animals Kienle et al. (2021); Lämmermann et al. (2013); Uderhardt et al. (2019); Poplimont et al. (2020); Isles et al. (2021); Chtanova et al. (2008); Khazen et al. (2022); Ng et al. (2011), the *in vivo* context presents several challenges for mechanistic dissection of the swarming process. For example, multiple cell types potentially modulate swarm initiation and propagation Uderhardt et al. (2019), there are a multitude of diffusive signals following tissue damage Uderhardt et al. (2019); Lämmermann et al. (2013), and the *in vivo* migration environment is complex. These challenges make it difficult to mechanistically dissect the regulation of swarming. Furthermore, a focus on model organisms precludes the analysis of swarming for human neutrophils despite known differences in primary human neutrophil behaviors compared to model systems Nauseef (2023); Siwicki and Kubes (2023). To address these limitations, we are leveraging an *ex vivo* assay for studying human neutrophil swarming in response to controllable, reproducible, well-defined cues. This assay expands upon a previously-developed *ex vivo* swarming system Reátegui et al. (2017); Hopke et al. (2020) in which a defined grid of heat-killed *Candida albicans* “targets” are spotted on a slip of glass to act as an array of swarm initiation points. Healthy human primary neutrophils isolated from whole blood are added to the assay, and swarming responses are observed with live cell microscopy.

## Fast-moving multicellular Ca^2+^ waves define the zone of recruitment in human neutrophil swarming

To monitor cell-cell communication in conjunction with more traditional swarming readouts like motility, we used a cytosolic calcium dye. Neutrophils increase cytosolic calcium following exposure to primary chemoattractants as well as swarming cues such at LTB4 Molski et al. (1981), and we envisioned that monitoring calcium influx would enable us to follow the propagation of swarming signals and correlate these to the regulation of directed cell movement (**Figure 1a**). Furthermore, calcium signals at the core of injury sites *in vivo* have been shown to be linked to key swarm signaling events Poplimont et al. (2020), and the simpler planar environment and higher signal-to-noise imaging of our assay should enable a more sensitive detection of swarm signal propagation to neutrophils far from the heat-killed *Candida albicans* target. Calcium signaling has been studied during mice and zebrafish swarming Poplimont et al. (2020); Khazen et al. (2022) but not during human neutrophil swarming. Following neutrophil introduction to the assay, we observe a calcium influx of the neutrophils that are in contact with the fungal target as well as multiple multicellular waves of calcium activity that radially propagate away from the target to surrounding neutrophils (**Supplemental Movie SM1**). An example time course of these waves is shown in **Figure 1b** with the overlayed tracked boundary of the calcium wave. Wave tracks are calculated by fitting a circle to a cloud of cells that are determined to be active via calcium signal binning and grouped using the ARCOS algorithm Gagliardi et al. (2022) (**Supplemental Figure S1a,b; Supplemental Movie SM2**). Wave movement is roughly isotropic, and the fitting of waves to a circle enables the tracking of wave propagation kinetics (**Supplemental Figure S1c,d**). Immediately following the passage of a calcium wave, cells rapidly polarize and radially migrate towards the wave origin site. Cells within the wave perimeter move towards the fungal target, while cells outside the wave perimeter lack coordinated movement (**Figure 1c,d**). By analyzing cell responses across many targets and donors, we observe that most, if not all, movement occurs within the boundaries of the tracked calcium wave, indicating that these wave fronts represent an effective boundary of the swarming guidance cue reception (**Figure 1e**). Though the propagating calcium waves delimit the zone of recruited neutrophils, they operate at very different timescales with respect to neutrophil movement. Calcium waves move an order of magnitude faster than the resulting neutrophil chemotaxis towards the wave origin (**Figure 1f**), and calcium waves propagate in an opposite direction to cell movement during swarming.

**Figure 1.**
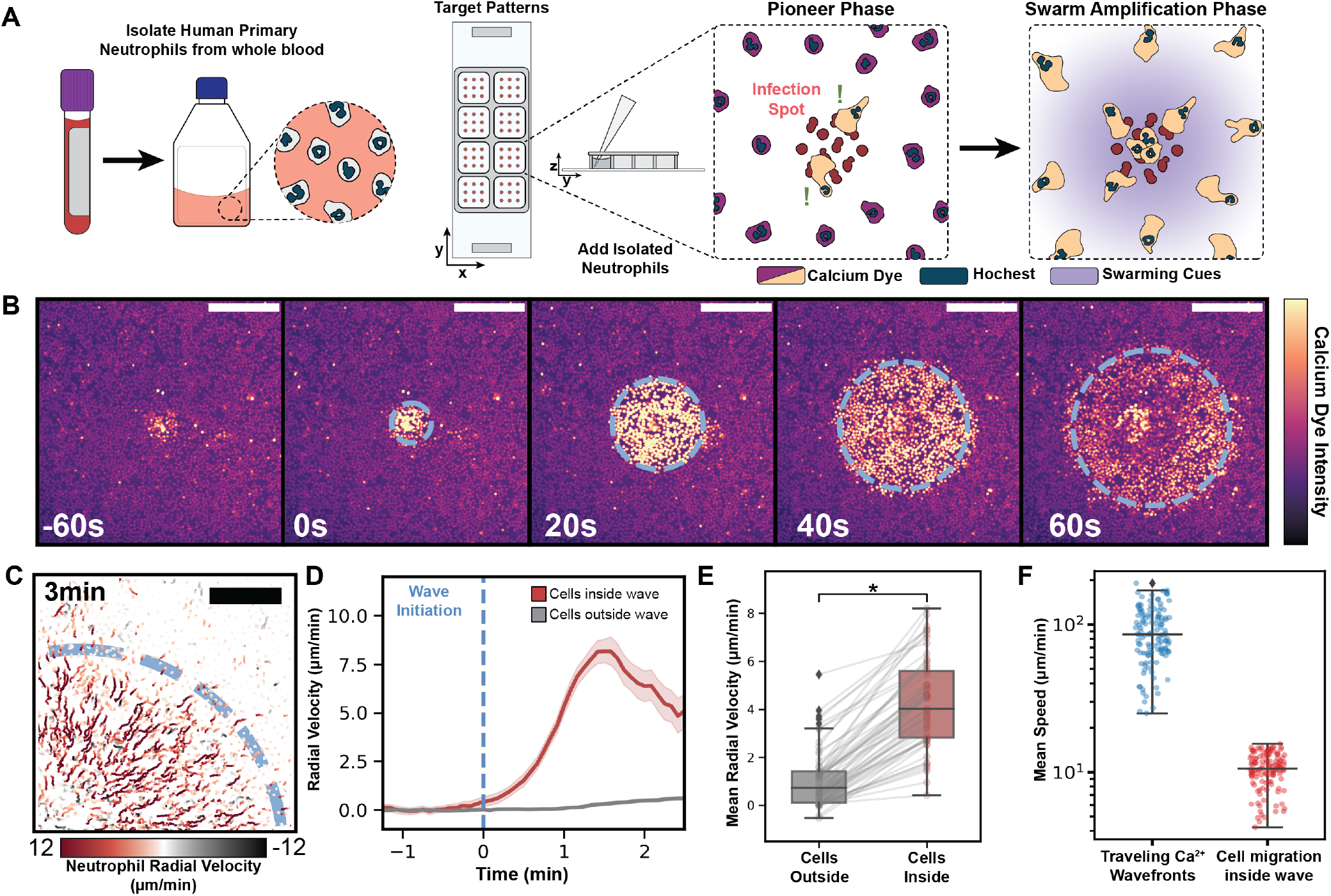
Fast-moving multicellular Ca2+ waves define the zone of recruitment in human neutrophil swarming. (**A**) *Ex vivo* assay for neutrophil swarming with self-amplified recruitment of cells in response to local site of heat-killed *Candida albicans*. Human neutrophils are isolated via immunomagnetic selection and dyed with CalBryte 520-AM (cytosolic Ca2+ reporter, used to monitor neutrophil detection of pathogen and swarming cues) and Hoechst 3334 (nuclear marker, used to track cell movement). Cells are placed on printed targets of heat-killed *Candida albicans*, and neutrophil swarming responses are monitored via confocal microscopy. (**B**) Time lapse sequence of a multicellular wave of neutrophil Ca2+ activity following detection of the fungal target. Dotted line represents wave boundary as tracked by our analysis software; scale bar = 200 μm. (**C**) Ca2+ wave defines the zone of neutrophil recruitment towards the fugal target. Final radius of Ca2+ wave from cells in 1b is shown in blue. Cell tracks for 3 minutes following wave initiation are indicated with color corresponding to radial velocity towards the wave center; scale bar = 100 μm. (**D**) Average radial velocity for cells inside versus outside the final wave boundary is plotted over 4 minutes for a population of cells during a single Ca2+ wave (Cell tracks inside the wave n = 3720, outside wave tracks n = 9499). Wave boundary predicts zone of recruitment during swarming. (**E**) Comparison of mean radial velocity for cells inside versus outside largest Ca2+ wave boundary in each target ROI. Tracks were averaged over the duration of the wave event, and lines connect mean radial velocities of outer versus inner cells for each target ROI. Dependent paired t-test p-value = 5.8 × 10−27 (Target ROIs n = 56) (**F**) Average velocity of Ca2+ waves propagated across the field of cells during swarming compared to the average cell velocity for the cells within these wave boundaries. Propagated Ca2+ waves are approximately one order of magnitude faster than cell movement during swarming.

## Wave propagation during neutrophil swarming is consistent with active relay but not core diffusion

LTB4, an inflammatory lipid of the leukotriene family, is thought to be one of the key secreted molecules that regulates neutrophil swarming Lämmermann et al. (2013); Reátegui et al. (2017). Neutrophils release LTB4 in response to various damage-associated pattern molecules and pathogen-associated pattern molecules Reátegui et al. (2017); Afonso et al. (2012); McDonald et al. (1994), including *Candida albicans* Fischer et al. (2021). To test whether LTB4 reception is required for the rapid long-range calcium waves in our *ex vivo* swarming assay, we blocked LTB4 reception with the LTB4 receptor antagonist BIIL315 Birke et al. (2001). Blocking LTB4 receptors inhibited the rapidly propagating long-range calcium waves that accompany swarming. In the absence of LTB4 reception, a much smaller range, slow-moving Ca^2+^ wave was observed (**Supplemental Figure S2b,c; Supplemental Movie SM3**). Quantitative analysis of wave propagation in both settings indicate that the rapid, long-range calcium waves that accompany swarming are dependent on LTB4 reception (**Figure 2a**).

**Figure 2.**
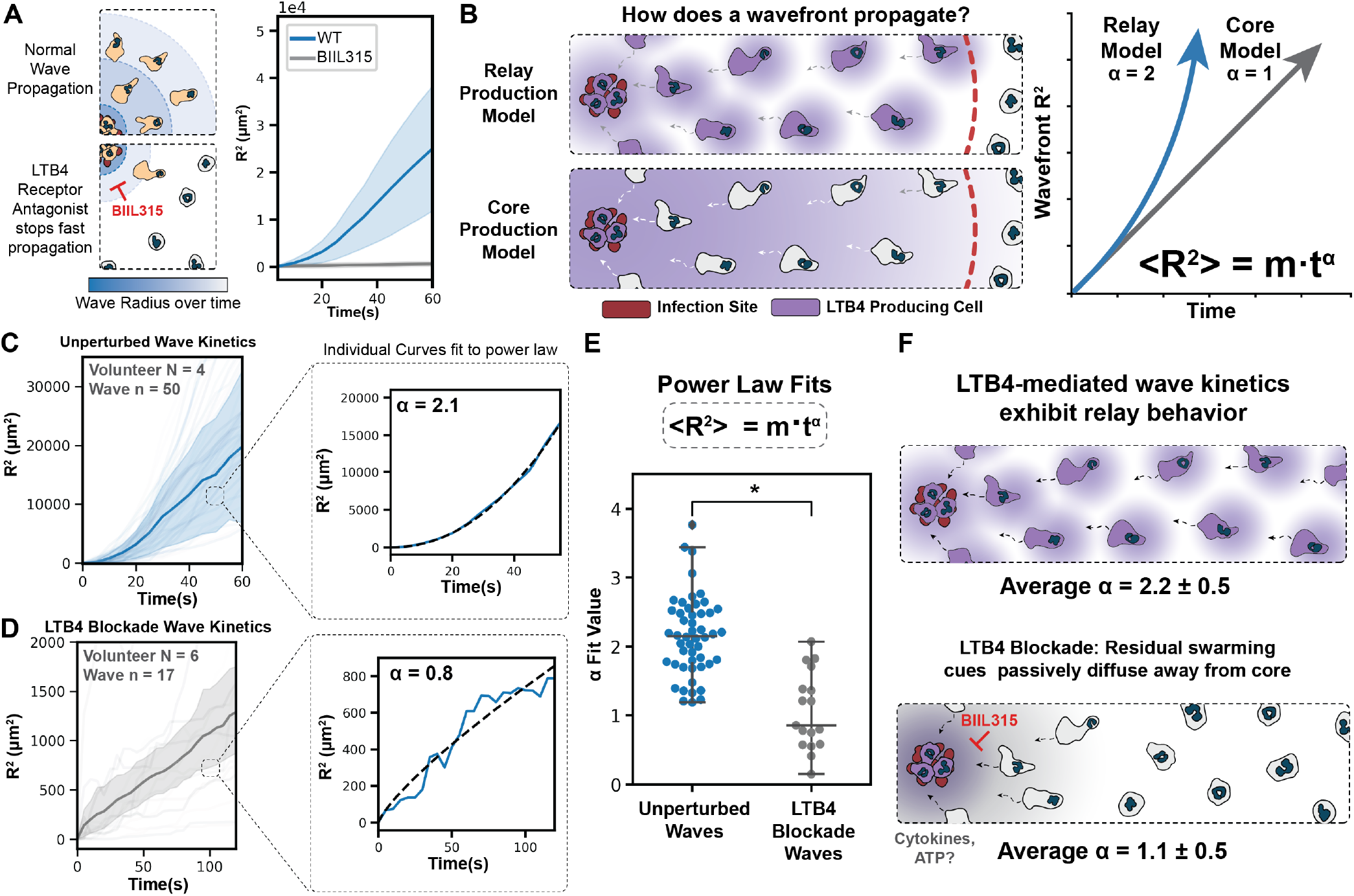
Wave propagation during human neutrophil swarming is consistent with active relay but not core diffusion. (**A**) Ca^2+^ wave propagation across a field of neutrophils during swarming for cells in the absence (50 waves) or presence (17 waves) of an LTB4 receptor inhibitor (1 μM BIIL315); averages shown with 95% confidence interval shaded. LTB4 reception is required for rapid, long-range Ca^2+^ wave propagation. (**B**) Two proposed models of LTB4 propagation from a site of infection to the rest of the field with the predicted kinetics of signal propagation shown for each. In a relay model of swarming Afonso et al. (2012); Lämmermann et al. (2013); Dieterle et al. (2019), each activated neutrophil releases LTB4, which stimulates the adjacent cell to release LTB4; this wave should spread with a constant velocity over time, giving an *α* of 2. For a core production model of swarming Poplimont et al. (2020), only neutrophils at the center of a swarm produce LTB4, which moves across the field through passive diffusion; this wave should spread with an *α* of 1 (linear on R^2^ plot). (**C**) Average early wave kinetics for control cells with mean and 95% confidence interval. Individual tracks are fit to a power law equation to determine the *α* of wave propagation for each experiment. (**D**) Wave propagation for LTB4R-inhibited cells averaged with 95% confidence interval in grey are individually fit to a power law equation for the first two minutes following swarm initiation. (**E**) Resulting curve fit alpha parameters for both unperturbed and LTB4R blockade conditions are plotted, with control cells showing wave propagation consistent with active relay. LTB4 reception-inhibited cells showed residual wave propagation consistent with core diffusion. Welch’s t-test P-value = 1.7 × 10^−7^ (**F**) Summary of this figure. LTB4-mediated waves propagate with kinetics consistent with active relay (and not core diffusion), while the zone of cell activation for LTB4-blockaded cells propagates away from the site of heat-killed *Candida albicans* with kinetics consistent with simple diffusion.

We next investigated how the LTB4 ligands are propagated from the target site to the rest of the field during swarming. Two models have been proposed for the signal amplification observed during swarming. For the relay model of swarming, pioneer neutrophils at the site of infection secrete LTB4, which activates surrounding neutrophils to secrete additional LTB4, thereby continuing the relay Lämmermann et al. (2013); Afonso et al. (2012); Dieterle et al. (2019). This active relay mechanism could enable neutrophils to collectively signal across significant distances from the injury/infection site, analogous to cell-cell signal propagation in aggregating Dictyostelium Shaffer (1975); van Oss et al. (1996). In contrast, for the core production model of swarming, only neutrophils in direct contact with the site of injury/infection produce significant LTB4, which then passively diffuses into the tissue to attract more neutrophils Poplimont et al. (2020). These two swarming models give very different predictions for the dynamics of propagation of the LTB4 wavefront as it moves away from the site of infection. The relay model predicts a wave that travels at a fixed velocity because of continuous signal re-amplification at the traveling front. This produces an activation zone whose area (wave fit circle radius squared / R^2^) scales quadratically in time. In contrast, since the core diffusion model predicts LTB4 production restricted to the central source, this system is limited by diffusion, and therefore its activation area (*R*^2^) scales linearly in time (**Figure 2b**). We tracked the radius^2^ versus time for multiple waves across many healthy donors in control cells (**Figure 2c**) versus LTB4-blockaded cells (**Figure 2d)** and fit a power law to each individual wave trajectory. This analysis reveals that control cells exhibit an LTB4 reception-dependent wave propagation with an alpha close to 2, consistent with the active relay model of swarming and inconsistent with the core production model of swarming (**Figure 2e**,**f**). In the absence of LTB4 signal reception, cells exhibit a slower, smaller pattern of signal propagation with an alpha close to 1, which is consistent with core diffusion. These data suggest that the residual release of non-LTB4 ligands from cells on the target passively diffuse to the rest of the field (**Figure 2e,f**). These two examples establish that we can distinguish between diffusive waves and relay-mediated ones, and LTB4-mediated waves exhibit relay-like ballistic spreading in the early phase of wave propagation.

## Swarming relay waves self-extinguish through an NADPH Oxidase negative feedback loop

Our observation that neutrophil swarming cues radiate from killed yeast targets via active relay might be expected to produce waves that continue to propagate as long as there are nearby receptive cells to continue the wave. This is the expected behavior of actively relayed systems including action potentials, mitotic waves, and Dictyostelium aggregation Gelens et al. (2014). We previously modeled an active relay model for swarming in which cells that pass a threshold amount of LTB4 themselves create more LTB4, as diagramed in **Figure 3a** Dieterle et al. (2019). Once initiated, these waves continue to propagate indefinitely given a field of responsive cells. In contrast with the predictions of a simple relay model, our experimentally-observed waves propagate and then abruptly stop (**Figure 3b,c**), indicating a more complex mode of regulation than positive feedback alone. While negative feedback loops could theoretically collaborate with positive feedback to generate self-extinguishing relay, we were only able to find one such example in the literature. In this work, a 1D model of two separately-generated waves of diffusing activator and inhibitor were able to generate a fixed radius of response in a cell-free system Ataullakhanov et al. (1998). We sought a simpler potential mechanism of wave stopping that does not depend on two independently propagated waves (see **Supplementary Modeling Text, Supplemental Figure S3**) and that is consistent with the dynamics of wave termination in our experimental context. This self-extinguishing relay model depends on two different responses initiated at two different concentrations of LTB4. Cells begin generating a non-diffusive internal inhibitor once they pass a low extracellular threshold of LTB4. After passing a higher threshold of extracellular LTB4, cells begin to release LTB4 themselves (**Figure 3d**). In this system, cells farther from an initiation event will have an increasing time delay between crossing the lower ‘inhibition circuit initiation’ threshold and the higher LTB4 production threshold. When a cell passes the threshold for inhibition, it begins accumulating the inhibitory cue that limits the relay strength of that cell. Close to the source of the activating cue at the target, only a small amount of inhibitor is generated prior to LTB4 production, and relay is still possible. Further from the source, the inhibitor accumulates for longer periods before the LTB4 production threshold is passed, thereby terminating further signal relay. This model can recapitulate the experimentally-observed wavefront dynamics (**Figure 3d**).

**Figure 3.**
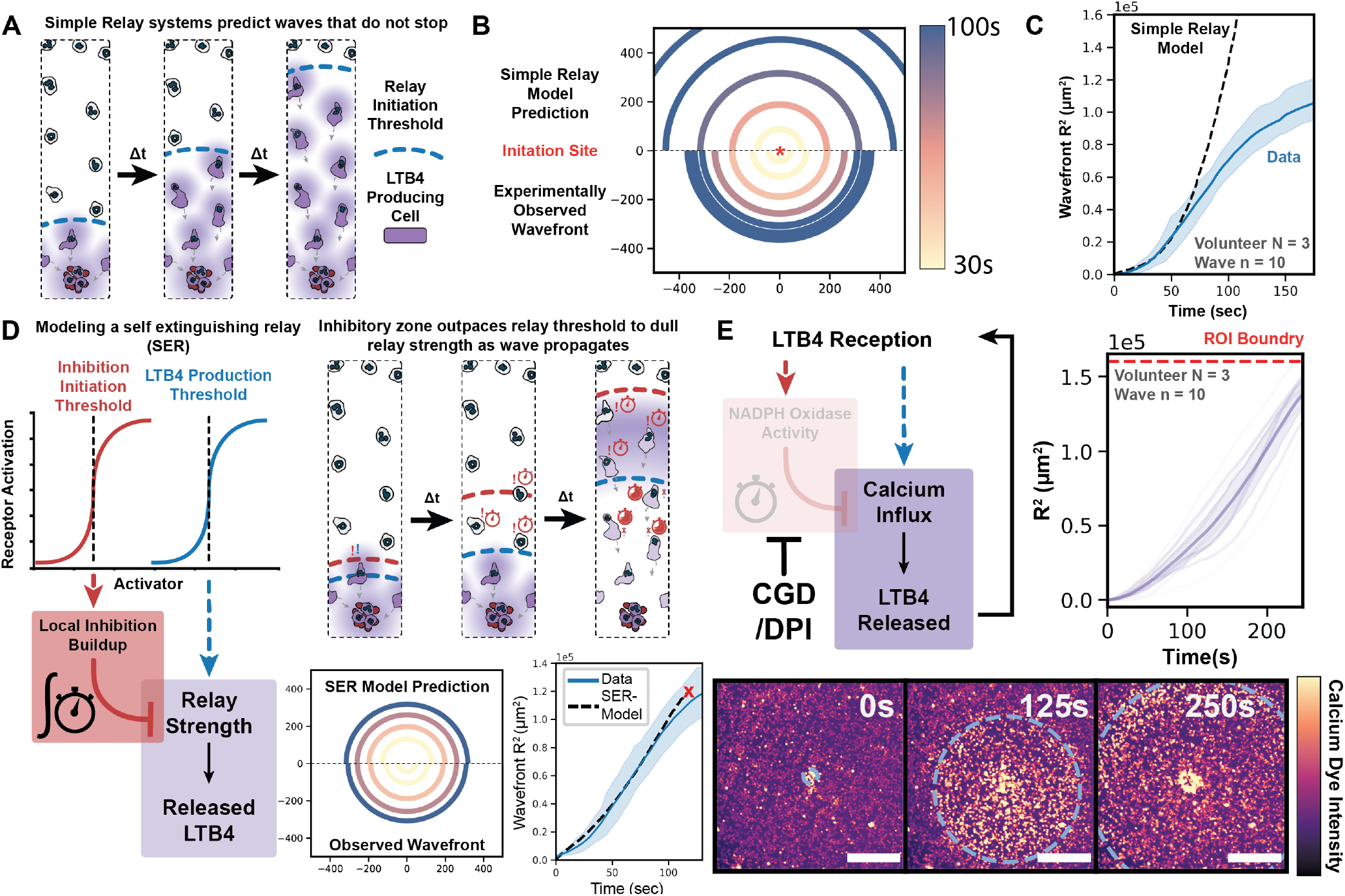
Neutrophils employ a NADPH-oxidase-based negative feedback loop to generate a self-extinguishing relay during swarming. (**A**) Previously-proposed simple diffusive relay model of swarming Dieterle et al. (2019). As cells exceed a threshold amount of extracellular LTB4, they begin to release more LTB4 to their surroundings. This wave propagates indefinitely once initiated in a homogenous field of cells. (**B**) Comparing the simple diffusive relay prediction with an average of experimentally-observed swarming wavefronts shows how wave behavior diverges, as experimental relay waves come to a stop, whereas simple relay propagates indefinitely (Axes in *μ*m. Wave n = 5. Volunteer N = 2). (**C**) While experimental waves initially propagate with a relay-like R^2^ relation, in the latter phase the wave decelerates then stops. (**D**) With the introduction of an activation-dependent inhibitor, it is possible to generate models of a self-extinguishing relay. Here different extracellular LTB4 thresholds for production of a local inhibitor (low threshold) versus release of more LTB4 (high threshold) generate an area of local inhibition that limits the wave propagation. Since the local inhibition requires accumulation over time to inhibit relay, wave termination is only achieved at further distances from the source. Our self-extinguishing relay model yields wavefront kinetics that closely resemble the experimentally observed Ca^2+^ swarming wavefront. The red X denotes where the model predicts the wave extinguishing (2D panel axes in *μ*m.). (**E**) Here we test a candidate activation-dependent inhibitor for the self-extinguishing relay during swarming. In neutrophils, LTB4-stimulated activation of the NADPH Oxidase complex limits the Ca^2+^ influx that potentiates LTB4 synthesis Song et al. (2020). Inhibition of NADPH oxidase activity via the NADPH oxidase inhibitor DPI produce swarming waves that no longer stop in the observable microscopic field (lower panels, scale: 200 μm) and mimic simple relay dynamics throughout their propagation (right panel) (Volunteer N = 3, Wave n = 11).

What might serve as the activation-dependent inhibitor in this self-extinguishing relay mechanism? The inhibitor should be produced downstream of LTB4 reception, and the accumulation of this inhibitor should attenuate LTB4 production. NADPH oxidase activation satisfies both conditions. NADPH oxidase is used in ROS-dependent pathogen killing downstream of LTB4 and other chemoattractants Song et al. (2020); Hopke et al. (2020), and the inhibition of NADPH oxidase activation potentiates LTB4 production and neutrophil accumulation during swarming Song et al. (2020); Hopke et al. (2020); Henrickson et al. (2018); Roxo-Junior and Simão (2014); Hamasaki et al. (1989). If NADPH oxidase activity is a critical component in self-extinguishing relay, we would expect NADPH oxidase inhibition to prevent swarming wave termination, thereby reverting the swarming process to a simple relay system (as in **Figure 3a**). Indeed, the NADPH oxidase inhibitor DPI (**Figure 3e, Supplemental Movie SM4**) abolished the ability of swarming waves to self-extinguish, with swarming waves continuing to propagate from the targets out of the microscopic field of view. To ensure that this effect is not an off-target of DPI, we analyzed neutrophils from Chronic Granulomatosis Disease (CGD) patients with a genetic defect in neutrophil NADPH oxidase machinery. CGD neutrophils exhibit similar non-extinguishing wave dynamics as DPI-treated healthy donor cells (**Supplemental Figure S4a**). While both DPI based NADPH Oxidase-inhibited and a CGD donor wave activity led to strong recruitment of the cells within a calcium event, they only generated a single large wave compared to the multiple waves observed in unperturbed cells (**Supplemental Figure S4b; Supplemental Movie SM4**).

## Neutrophils tune the size and number of swarming waves for homeostatic recruitment to targets

To investigate the physiological significance of multiple self-extinguishing waves (control neutrophils) versus a single uncontrolled relay wave (DPI-treated or CGD neutrophils), we next sought to investigate the relation between wave size/number and the resulting cell movement and recruitment. We hypothesized that these wave parameters may be actively adjusted to ensure robust homeostatic control of neutrophil recruitment across a wide range of initial conditions. We created quantitative metrics for both the propagating Ca^2+^ wave as well as neutrophil chemotaxis response that could be extended to multiple wave sizes and intensities. First, we aligned single-cell tracks for neutrophils entering the zone of recruitment and averaged their radial migration velocity profile for single, well-separated waves (**Figure 4a, Supplemental Figure S5a**). Upon entering a wave, neutrophils execute a discrete ‘run’ of movement towards the wave center but then revert to random, slower movement. Such a ‘step’ of movement inwards is consistent with the expected behavior from our self-extinguishing model where the chemotactic gradient decays behind the wavefront as the relay strength becomes more dominated by the local inhibitory mechanism. These data indicate that a single wave event does not recruit all cells to the center in one burst. To expand our analysis to multiple waves, we integrated the radial movement of all cells within a wave event (**Figure 4b, Supplemental Figure S5b**). Each wave independently recruited of a bolus of neutrophils toward a given target, with larger wave events more potently stimulating cell recruitment than smaller wave events (**Supplemental Movie SM5**). When looking across many experiments and donors, the total integrated wave area highly correlated with the Integrated Radial Movement of the cells within wave events (**Figure 4c**).

**Figure 4.**
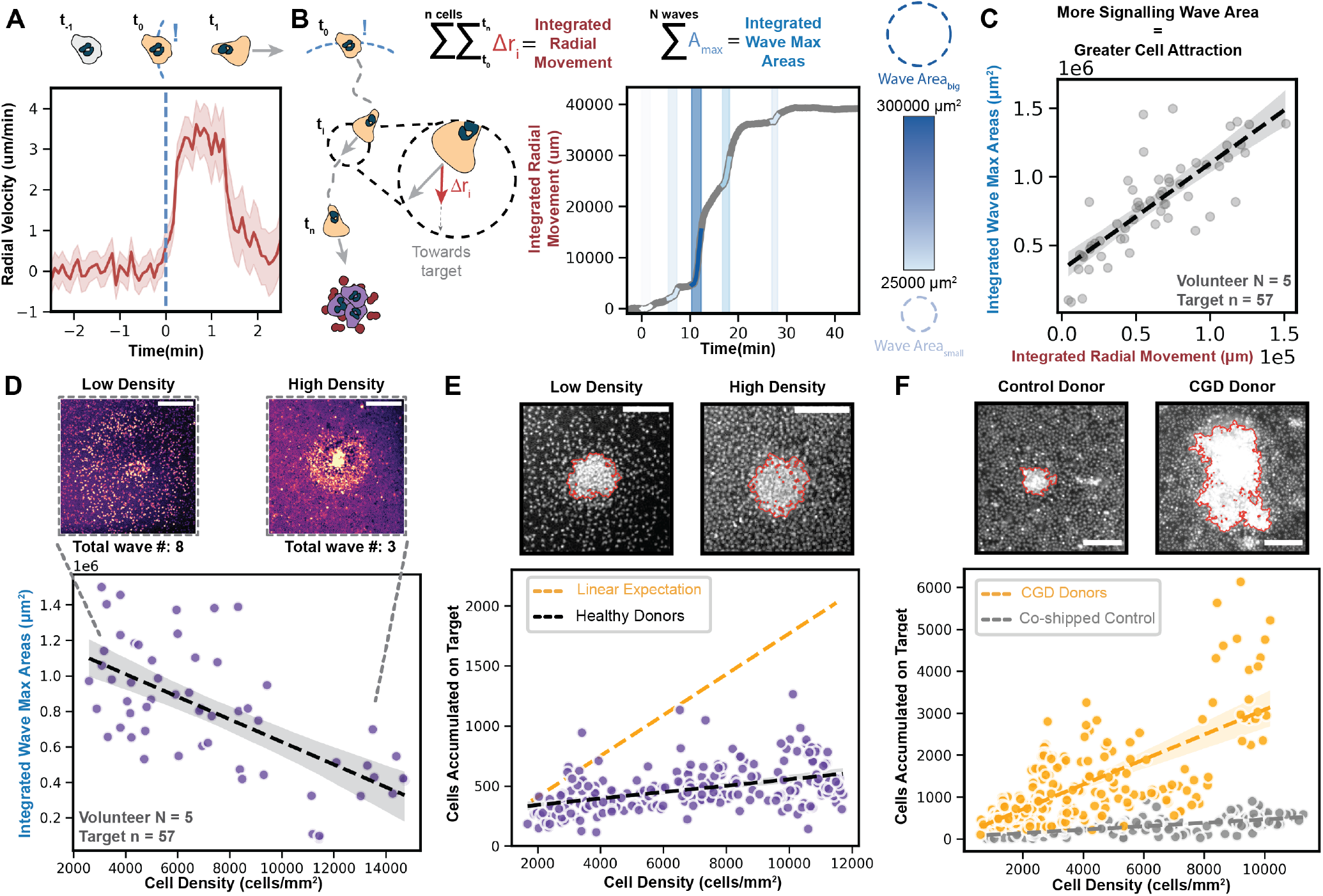
Neutrophils achieve homeostatic recruitment to sites of heat-killed *Candida albicans* by modulating the size and frequency of swarming waves. (**A**) Averaged radial velocities of neutrophil movement towards a target following time of exposure to swarming Ca^2+^ wave with 95% confidence interval plotted for 1758 cell tracks. A single wave produces a transient bolus of neutrophil recruitment. (**B**) All cellular tracks inside a wave are integrated for their radial movement (Cell track n = 5813); multiple Ca^2+^ wave events for the same target are plotted as shaded areas colored by maximum final area for an example ROI. Each wave induces a bolus of neutrophil movement towards the target, with larger waves inducing larger neutrophil responses. (**C**) Integrated radial movement of all cell tracks and integrated wave max area measurements calculated across 57 target ROIs demonstrate the strong positive correlation between Ca^2+^ wave signaling area and movement towards a target. Linear fit slope: 7.8 μm *±* 0.8, 95% confidence interval. A control where all cell tracks are considered is in Supplemental Figure S5C. (**D**) Relation of seeding density to cell accumulation at target. Integrated wave max area measurements taken for each target ROI over 45min. Linear fit slope: −6.3 × 10^4^ μm^2^ per 1000 cells/*mm*^2^ *±* 1 × 10^4^, 95% confidence interval. Lower cell densities elicit larger and more numerous swarming waves. (Scale bar: 200 μm). (**E**) Z-stack measurements taken across multiple ROIs per well 60 min after seeding swarming assays at different cell densities. Accumulation measured as integrated fluorescence at target site divided by median single-cell nuclear fluorescence. Low Density ROI Estimation: 402 cells, High Density ROI Estimation: 551 cells. Linear fit slope: 27 cells accumulate at target site per 1000 cells/*mm*^2^ *±* 4 cells; 95% confidence interval. Only a small change in final neutrophil target accumulation is seen over a 6x range of cell swarming densities, compared to expectation for swarming directly proportional to cell density (linear scaling, orange line) (Scale bar: 100 μm). (**F**) Widefield measurements taken across multiple ROIs per well 60min after seeding swarming assays with different cell densities of either control donor cells or CGD (NADPH oxidase-defective) human donor cells. Accumulation measured similar to E, see Methods. Control ROI Estimation: 295 cells, CGD ROI Estimation: 4780 cells. CGD Linear fit slope: 297 cells accumulate at a target per 1000 cells/*mm*^2^ *±* 19 cells, 95% confidence interval (Volunteer N=4, Target n = 253). Healthy control linear fit slope: 40 cells accumulate at a target per 1000 cells/*mm*^2^ *±* 4 cells (Volunteer N = 3, Target n = 157)(Scale bar: 100 μm).

We asked next whether cells tune the number or size of swarming waves for robust homeostatic control of swarming over a range of initial conditions. Towards this end, we seeded our swarming assay at a range of initial neutrophil densities. As seeding density increased, neutrophils generated smaller and fewer swarming waves, resulting in a decreased overall integrated calcium area (**Figure 4d, Supplemental Figure S5d, e**). This reduction in swarming waves with increasing cell density could potentially enable a homeostatic recruitment of the same number of neutrophils across a range of initial cell densities. To investigate this possibility, we monitored neutrophil recruitment to sites of heat-killed *Candida albicans* at a range of seeding densities over 60 min (**Supplemental Figure S5f, g; Supplemental Movie SM6**). Over a 6-fold change in initial cell density, there was a nearly constant accumulation of neutrophils at the target, with targets at the lowest densities attracting approximately 85% of the cells attracted at the highest cell densities (compared to 6x differences that would be expected for a non-homeostatic system) (**Figure 4e, Supplemental Figure S5h**). Because CGD cells are defective in the negative feedback loop that constrains wave size, we predicted that these cells should exhibit a defective homeostat and therefore recruit cells proportional to their surrounding density. Indeed, CGD cells exhibit a much stronger density dependence on recruitment than healthy donor cells (**Figure 4f, Supplemental Figure S4e, f**).

## Discussion

By leveraging a cell-cell signaling readout in a highly controllable *ex vivo* swarming assay, our work reveals self-extinguishing waves of human neutrophil recruitment to sites of heat-killed *Candida albicans*. The active relay is based on an LTB4-positive feedback loop (**Figure 2**), and the ability to self-extinguish depends on LTB4-based NADPH-oxidase activation (**Figure 3, 4**). The active relay enables cells to transmit guidance cues much farther and faster than would be possible if only the cells directly on the target secreted swarming signals. Furthermore, the propagated waves may generate temporally-evolving guidance cues that enable more effective chemotaxis than static spatial gradients Geiger et al. (2003); Tweedy et al. (2016); Aranyosi et al. (2015); Skoge et al. (2014). Behind a wavefront, decaying gradients could further prevent cells from over accumulating at the core of swarms as cellular memory mechanisms lose strength Skoge et al. (2014) and as chemotaxis-inhibitory LTB4 metabolites accumulate Archambault et al. (2019); Pettipher et al. (1993); Tweedy et al. (2016). The self-extinguishing nature of the relay enables the system to adjust the number and size of swarming waves to achieve robust homeostatic control of neutrophil recruitment over a wide range of initial cell concentrations. Disruption of this homeostat has severe consequences for neutrophil recruitment *in vivo*. Previous work has shown that CGD patient neutrophils (that are defective in NADPH-oxidase activation) overproduce LTB4 and hyperaccumulate at sites of injury/infection *in vitro* and *in vivo* Song et al. (2020); Hamasaki et al. (1989); Henrickson et al. (2018); Hopke et al. (2020); Dinauer (2019). Our work shows that these cells are also defective in the negative feedback arm that limits the range of swarming signals and therefore lack the homeostat that normally constrains cell swarming.

Our *ex vivo* system demonstrates that human neutrophils can modulate their cell-cell signaling to limit the range of cell swarming without feedback from other cell types or environmental cues. In future work, it will be interesting to probe how other cell types influence the initiation, termination, and resolving phases of swarming. We have focused on killed yeast-mediated neutrophil swarming (whose closest parallel is likely an infected lymph node Chtanova et al. (2008); Lämmermann et al. (2013)), but it will be interesting to compare how sterile injury Lämmermann et al. (2013); Ng et al. (2011); Poplimont et al. (2020); Park et al. (2018) and combined injury/infection Poplimont et al. (2020); Chtanova et al. (2008) change the dynamics of the swarming process. Other swarming terminators (such as Grk2 activation Kienle et al. (2021)) control the duration of swarming, whereas the NADPH oxidase negative feed-back circuit studied in our work also controls the spatial range of swarming signal propagation. It will be interesting to probe how these swarming termination programs relate to one another. Finally, as there are a number of other human diseases with known swarming defects Knooihuizen et al. (2021); Alexander et al. (2021); Barros et al. (2021), it will be interesting to determine how these disease states interact with our model parameters and effect the relay system we study in this work.

## Methods

For a detailed description of all the methods and code used in this work, please see Supplemental Methods.

## Supporting information

Supplemental Theory Text

## Acknowledgements

We thank members of the Weiner, Amir, and Irimia lab for helpful discussions, particularly Nick Martin and Henry De Belly for a critical reading of the manuscript. We thank Paul Dieterle, Wencheng Ji, and Allyson Sgro for their detailed feedback and advice regarding the theoretical model. We also thank the blood donors and phlebotomists without whom this project would have been impossible. This work was supported by an AHA predoctoral fellowship (ES), the National Institutes of Health grants (GM118167 to ODW, GM092804 to DI, R01 AI132638 to MKM), the National Science Foundation/Biotechnology and Biological Sciences Research Council grant (2019598 to ODW), the National Science Foundation Center for Cellular Construction Grant (DBI-1548297 to ODW), the Department of Defense (HU00012020011 to MKM) the Novo Nordisk Foundation grant for the Center for Geometrically Engineered Cellular Systems (NNF17OC0028176 to ODW), the Clore Center for Biological Physics (AA), the NSF CAREER Grant (1752024 to AA), and the Shriners Hospitals for Children (71010-BOS-22 to DI). CZ was supported by the Intramural Research Program of the National Institutes of Health Clinical Center and the National Institute of Allergy and Infectious Diseases. Finally, we would like to thank the Center for Advanced Light Microscopy-Nikon Imaging Center at UCSF for their support in using the CREST/C2 Confocal microscope. BIIL 315 was kindly provided by Boehringer Ingelheim via its open innovation platform opnMe, available at https://opnme.com.

## Supplementary Figures

**Figure S1.**
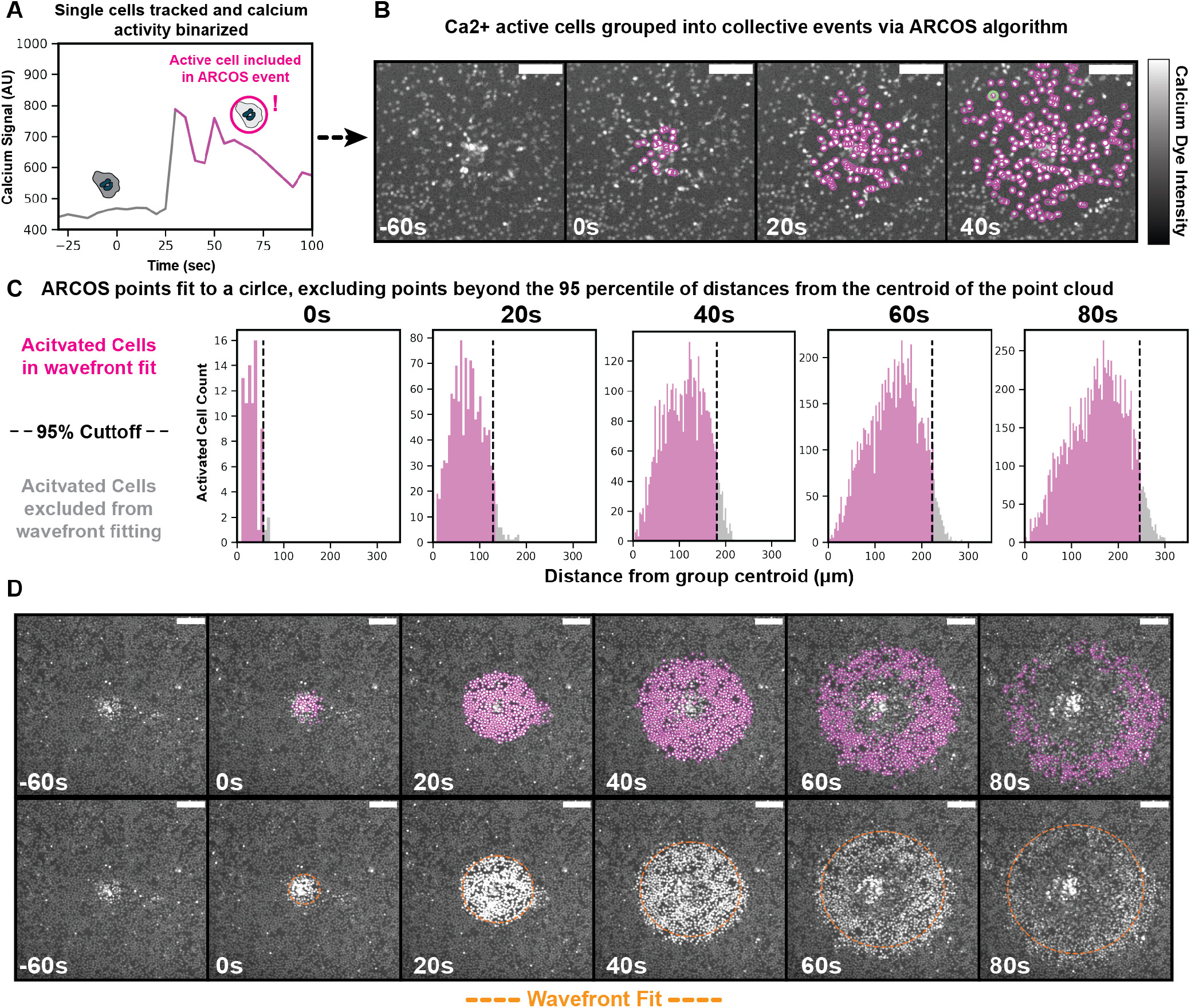
Tracking single cell Ca^2+^ behavior to quantify swarm wave propagation. (**A**) Single cell tracks are created using trackpy to link together frames of nuclei identified by a custom trained StarDist model. Over the course of each cell track, calcium signals are collected and binned based on a manually-set percentage threshold; normally an instantaneous reading 15% above the track mean is considered ‘on’. (**B**) Calcium ‘on’ cells are then grouped in space and time using the ARCOS algorithm to denote groups of spatiotemporally coordinated ‘on’ cells. A human-in-the-loop strategy is employed to ensure that waves are optimally grouped in each image ROI based on the tuning two parameters of neighborhood grouping size and calcium binning threshold. Minimum cluster size and minimum collective event duration values are held fixed for all analyses. Tuned values are recorded for each ROI used in this work and stored with the data files; scale bar = 100 μm. (**C**) To analyze how these ARCOS-grouped events spread in time, we fit the events to a circle of minimum diameter. To reduce fitting noise, we take the ARCOS-grouped point cloud centroid mean for each timepoint and then exclude the points in the top 5% percentile of distances to this centroid. This process enables us to fit a circle to each Calcium wave for later analysis. (**D**) Each ARCOS-grouped point cloud is shown in the top panels for the corresponding ROI for **Figure 1b**. The circle of best fit is shown below each point cloud; scale bar 100 μm.

**Figure S2.**
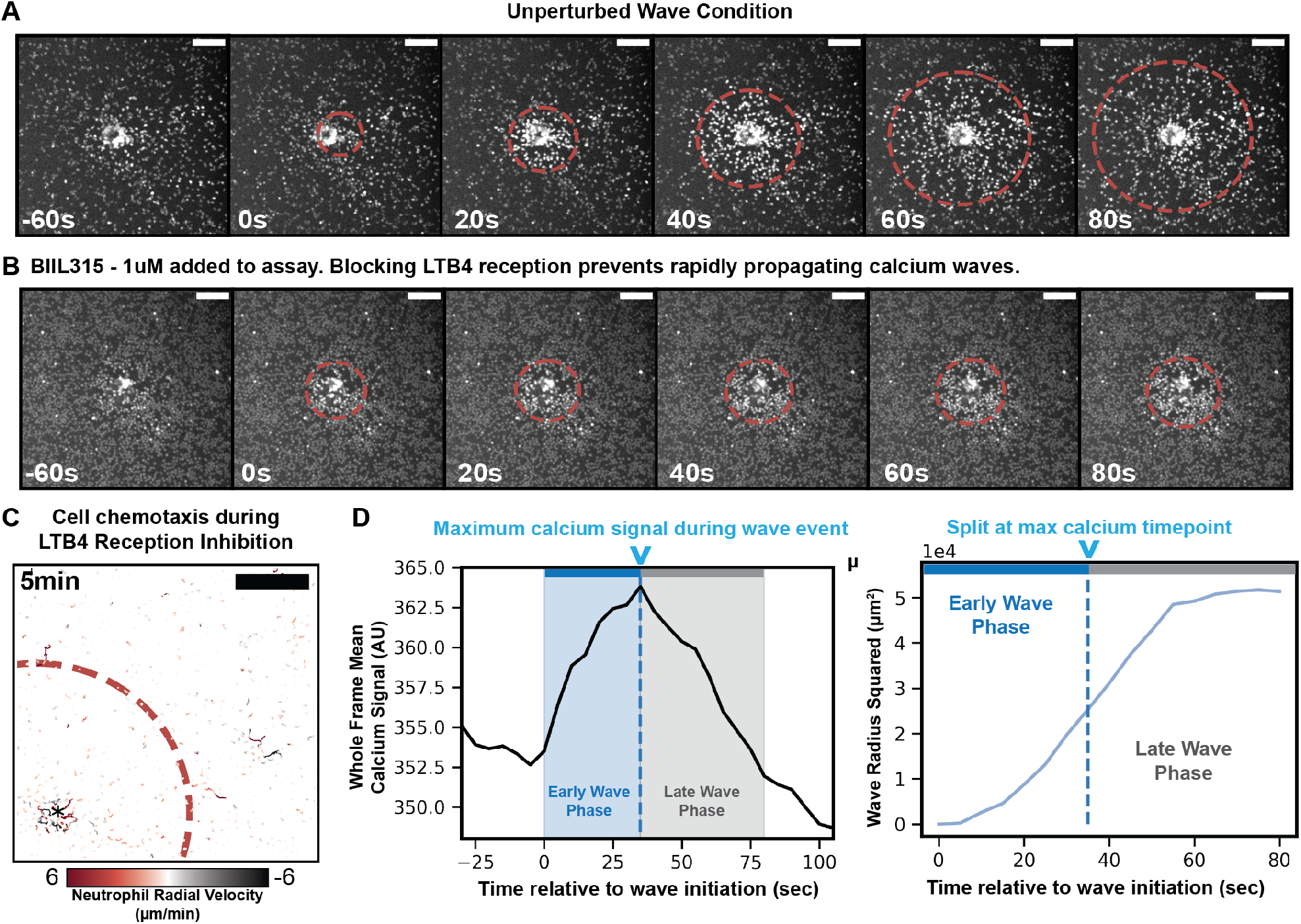
Tracking unperturbed and LTB4 reception inhibited Ca^2+^ waves; defining early and late wave phases. (**A**) Ca^2+^ wave propagation across a field of neutrophils during swarming in the absence (**A**) or presence (**B**) of an LTB4 receptor inhibitor (1 μM BIIL315); scale bar 100 μm. (**C**) Cumulative cell chemotaxis 5 minutes after calcium wave initiation in the presence of 1 μM BIIL315 (LTB4R Antagonist). Cell movement towards the target is observed for a much smaller subset of cells within the boundary of the Calcium wave compared to unperturbed cells (compare to **Figure 1c**); scale bar 100 μm. (**D**) To study early wave kinetics in our unperturbed conditions, we segmented waves into two phases – ‘early’ and ‘late’. We define early waves (blue region to left of v) as the period of calcium signal increase integrated across the wave, whereas late waves (grey region to right of v) correspond to the period of calcium signal decrease integrated across the wave. A single representative wave segmentation is shown for the calcium channel and also for the corresponding positions in the wave radius squared plot.

**Figure S3.**
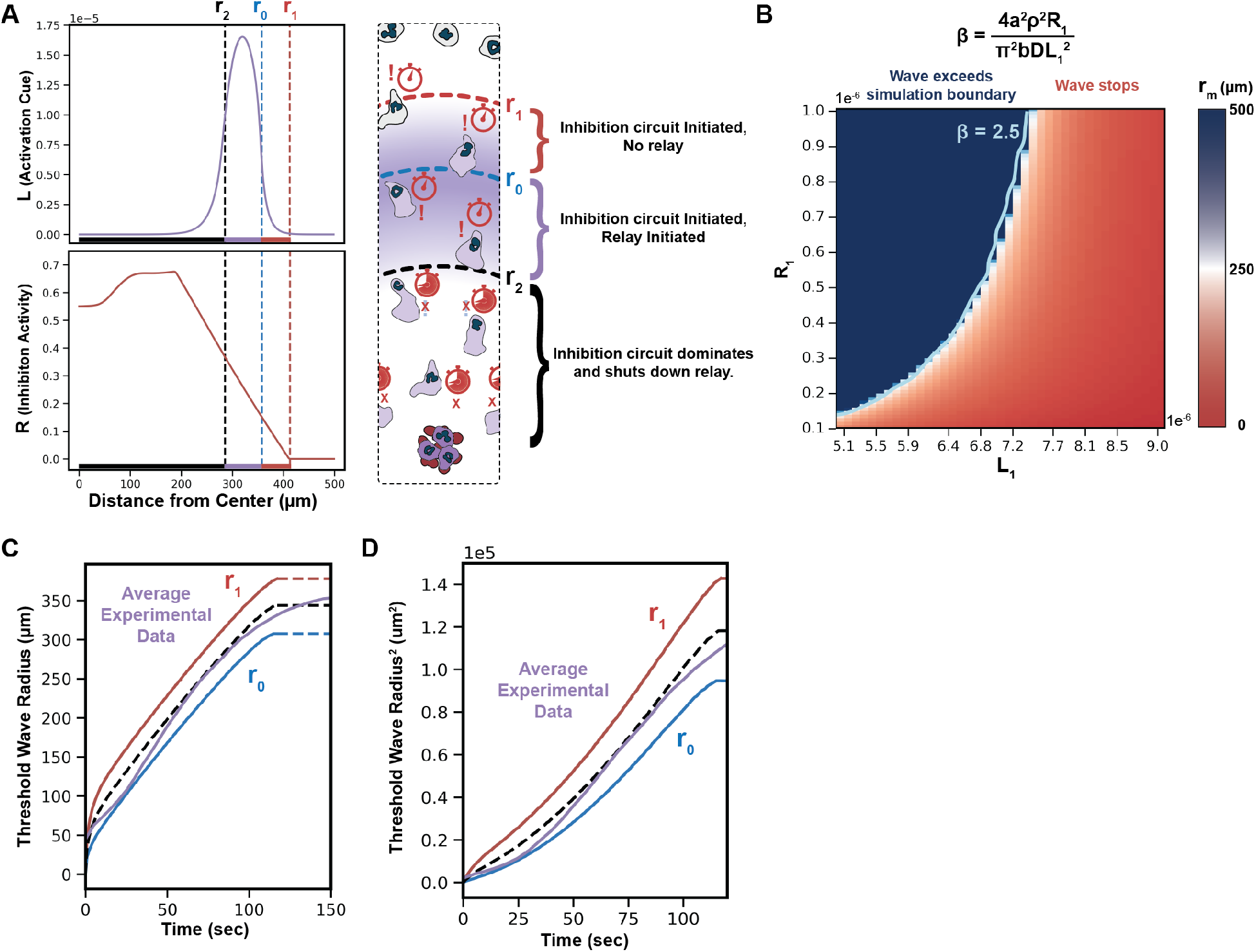
Modeling a self-extinguishing relay system. (**A**) Illustration of our model with a simulation snapshot. The simulation is generated by **Eq. 2** (**Supplemental Theory Text**) with parameters as follows (all spatial units are ms and all temporal units are seconds): *D* = 125, *ρ*= 10*−*2, a= 10*−*2, *γ*_*L*_= 2 × 10^−3^, *γ*_*R*_= 1 × 10^−3^, *L*_1_= 7.4 × 10^−6^, *L*_2_= 0.01L_1_,R_1_*/b*= 37, *m* = *n* = 10, and *k* = 20. Simulations are confined within r=500 circular polar coordinates with absorbing Dirichlet boundary conditions. The initial source is a constant source at *r <* 10 for *t <* 50 with amplitude *s*_*i*_ = 10*−*3. The activator production is only possible in (*r*_2_, *r*_0_), the area in which activator concentration is above *L*_1_ and inhibitor concentration is below *R*_1_. (**B**) Test of the robustness and sensitivity to model parameters. By modulating *L*_1_ and *R*_1_, we find that the characteristic value of *β* from **Eq. 7** (**Supplemental Theory Text**) stands as a reliable predictor for the persistence of wave propagation. Persistent relay process is possible when *β >* 2.5 in simulations, as their maximum propagation distance *r*_*m*_ easily reaches our simulation limit (500 μm). Other parameters used for solving **Eq. 2** numerically in this parameter screen of *L*_1_ and *R*_1_ (units in *μ*ms and seconds): *D* = 125, *ρ*= 10*−*2, a= 10*−*2, *γ*_*L*_= 2 × 10^−3^, *γ*_*R*_= 1 × 10^−3^, *L*_2_= 0.01L_1_,b=10^*−*8^, *m* = *n* = 3, and *k* = 5. The source at the origin which initiates the diffusive relay is a constant source at *r <* 10 for *t <* 50 with amplitude *s*_*i*_ = 10*−*3. (**C**) Comparison of our model with the Ca^2+^ wave spatiotemporal dynamics (averaged over multiple experiments). Shown are the inhibitor wavefront (the radius at which the inhibitor concentration is above *L*_2_(*r*_1_)) and the activator wavefront (where the activator concentration is above *L*_1_(*r*_0_). Colored dashed line segments indicate the wave stopping points. Both waves show the ballistic motion followed by an abrupt stop, as observed in the experiments.The black dashed line indicates the mean between these two thresholds where we would expect a calcium pulse to occur. This line represents the model fit in **Figure 3d**. Model parameters in a simulation using **Eq. 2** (units in *μ*ms and seconds): *D* = 125, *ρ*= 10*−*2, a= 10*−*2, *γ*_*L*_= 2 × 10^−3^, *γ*_*R*_= 1 × 10^−3^, *L*_1_= 7.4 × 10^−6^, *L*_2_ = 0.01*L*_1_, *R*_1_*/b*= 120, *m* = *n* = 3, and *k* = 5. (**D**) Radius squared values from C.

**Figure S4.**
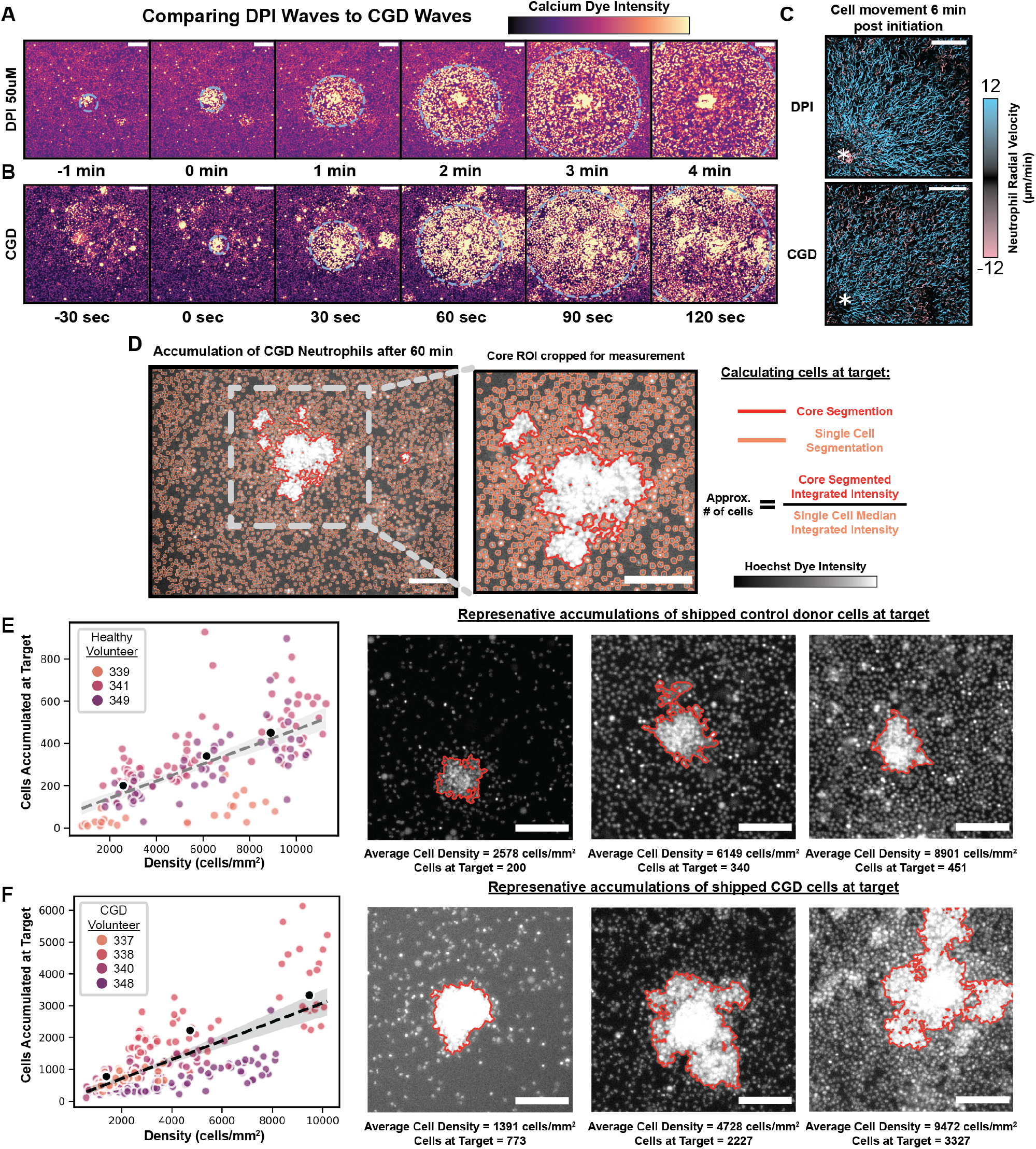
NADPH Oxidase inhibition results in uninhibited wave relay and a broken recruitment homeostat. Non-terminating calcium swarming waves two different modes of NADPH oxidase inhibition—the pharmacological inhibitor of NADPH oxidase DPI (**A**) and a human CGD donor defective in gp91 (Donor 338) (**B**). Calcium dynamics were visualized for cells exposed to targets of heat-killed Candida albicans (Scale bar: 100 μm). (**C**) The resulting cell movements are plotted for the six minutes following wave initiation. Cellular radial movement is both strong and persistent after a cell enters a CGD or DPI wave (Scale bar: 100 μm). (**D**) To measure cell accumulation at the target site one hour after seeding cells, single Hoechst fluorescence ROIs are taken using widefield microscopy. Each ROI is run through an algorithm (see Code) to identify single nuclei and clusters of cells. The larger ROI is used to calculate the average surrounding density of a target, whereas a smaller ROI is limited to the Candida target site, as determined from a DIC image taken before layering the neutrophils in each well. The total integrated fluorescence of an accumulation core is divided by the median-single-cell-integrated intensity in each ROI to measure the number of cells contained in a cluster (Scale bar: 200 μm). (**E**) Control healthy donor accumulation across varied cell density after 60min as measured by single-plane widefield Hoechst measurement. Linear fit slope: 40 cells accumulate at a target per 1000 cells/*mm*^2^ *±* 4 cells (Target n = 157). Cells potentially have a lower accumulation at low density because blood was shipped overnight from the NIH, leading to reduced initiation capability. This shipping effect is strongest with volunteer 339. Sites recorded to have 0 accumulation were excluded from the analysis (Scale bar: 100 μm). (**F**) CGD donor cell accumulation at a target 60 min after cell seeding is shown for each individual CGD patient; Linear fit slope: 297 cells accumulate at a target per 1000 cells/*mm*^2^ *±* 19 cells, 95% confidence (Target n = 253). Representative accumulations of CGD cells are shown with their corresponding average surrounding densities (Scale bar: 100 μm).

**Figure S5.**
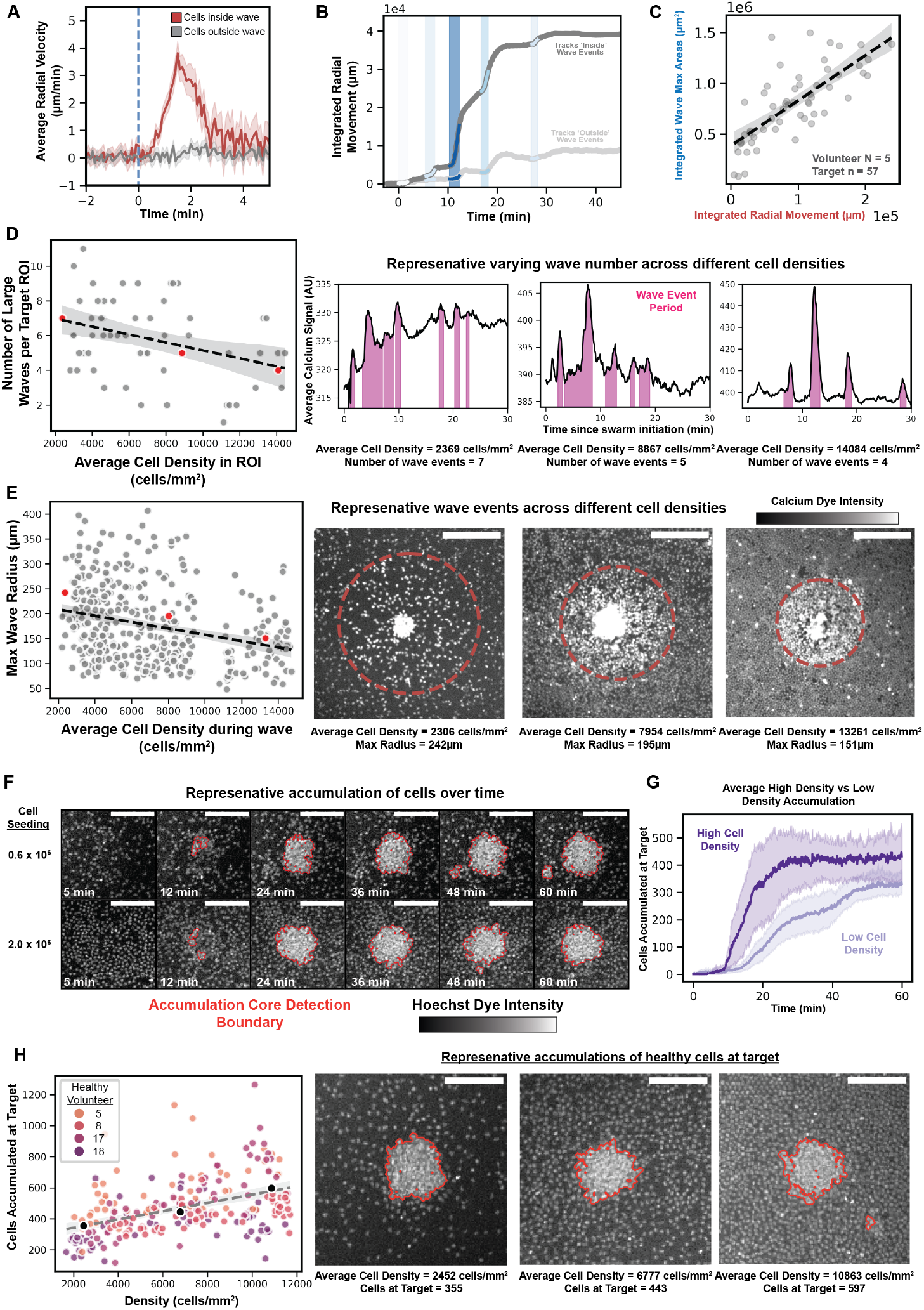
Neutrophils adjust swarming wave size and number to homeostaticaly control recruitment. (**A**) Averaged radial velocity of cells inside and outside of the wave used to align tracks in Figure 4A. Cell movement towards target is transient if only one wave passes through a given field of cells; (Tracks inside n = 3720, outside n = 9907); Confidence Interval: 95%. (**B**) The integrated movement towards a target for cells inside vs. outside a representative wave event; (Tracks inside n = 5096, outside n = 22105). (**C**) Integrated radial movement of all cell tracks both inside and outside a wave are calculated and compared with the summed wave areas in 57 target ROIs. This relation matches that in **Figure 4c** in which only cells inside a wave are included in the integrated radial movement measurement; Linear fit slope: 4.4 μm *±* 0.5, 95% confidence interval. (**D**) The number of waves emanating from a given target are inversely correlated with the seeding density of cells surrounding the target. Linear fit slope: -0.2 waves per 1000 cells/*mm*^2^ *±* 0.07, 95% confidence interval. Representative ROIs are shown with their corresponding max radius and measured cell density. (**E**) The maximum radius of waves emanating from a given target are also inversely correlated with the seeding density of cells surrounding a target. Linear fit slope: -6.4 *μ*m per 1000 cells/*mm*^2^ *±* 1.0, 95% confidence Representative ROIs are shown with their corresponding max radius and measured cell density (Scale bar: 200 μm). (**F**) As neutrophils begin to swarm around a given target ROI, their accumulation is monitored by thresholding the entire ROI and selecting for clusters of Hoechst florescence above a set area threshold. These clusters can be used to monitor cell accumulation at a target over time for different starting cell densities. Low and high seeding density examples are plotted during a 1 hr experiment (Scale bar: 100 μm). (**G**) Multiple targets at different cell seeding densities (High n = 7, Low n = 13) are monitored for cell accumulation over time; confidence interval 95%. Targets with a low cell density more slowly accumulate cells but continue to increase for a longer period, thereby approaching the same number of cells at the target as high-density seeding conditions. (**H**) End point accumulation measurements via confocal z-stack 60min post cell seeding in target wells; Linear fit slope: 27 cells accumulate at target site per 1000 cells/*mm*^2^ *±* 4 cells, confidence interval 95% (Volunteer N = 4, Target n = 230). Same data as **Figure 4e** broken out into volunteers, with representative points plotted in black. Representative greyscale images given of Hoechst intensity for in-focus z plane of an ROI with the accumulation boundary indicated. Sites recorded to have 0 accumulation were excluded from the analysis. (Scale bar: 100 μm)

## Supplementary Videos

**Figure SM1.**
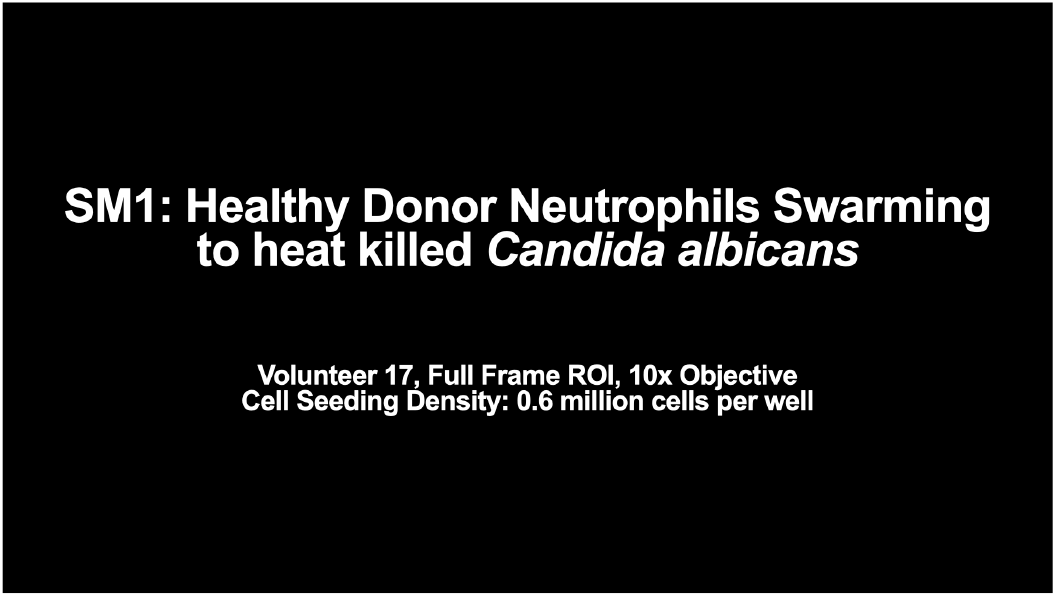
Healthy human donor neutrophils exhibit multiple pulsatile self-terminating swarming waves. Recording is initiated when human neutrophils are seeded on to targets consisting of heat-killed Caninda albicans. Calcium and Hoechst channels are imaged simultaneously on the Ti2 Crest confocal system described in Methods. Full frame (4 targets) and zoomed ROIs (one target) are displayed in the same video. Target well seeded at 600,000 cells in the well, Volunteer 17. Imaging media is RPMI with 0.4% HSA added and lacking phenol red.

**Figure SM2.**
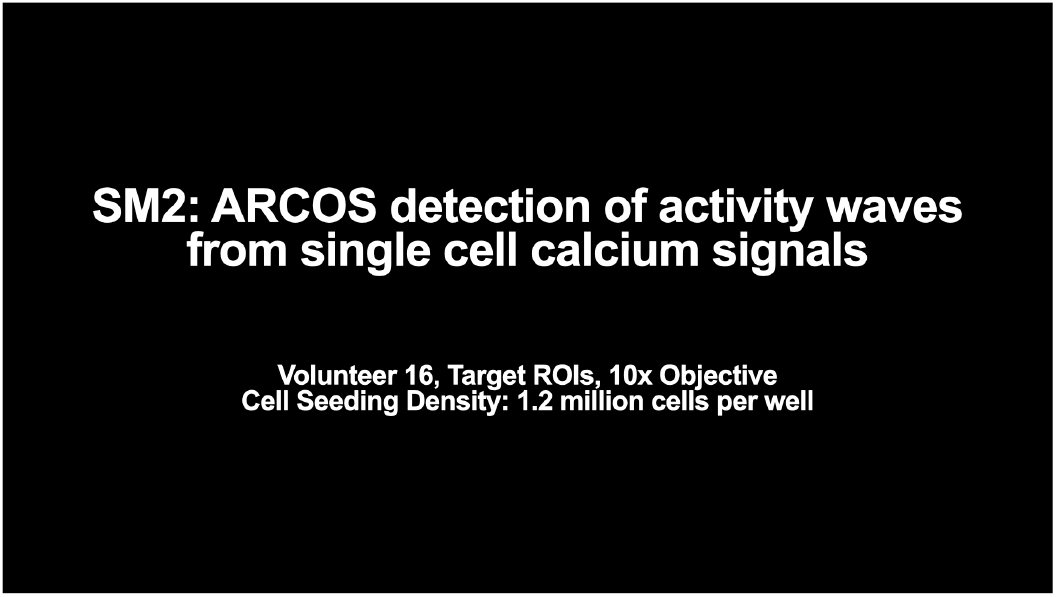
Example of automatic wave detection using ARCOS and designation of circular wave boundary. Same imaging conditions to SM1, target ROI displayed. Target well seeded at 1.2 million cells in the well, Volunteer 16. Imaging media is RPMI with 0.4% HSA and without phenol red. First panel shows a single cell tracked for its calcium behavior. When the circle drawn around the cell is yellow, the cell is considered off, and when its signal tracks 15% above the total signal mean, it is relabeled in magenta as ‘on’. The graph shown to the right corresponds to the example single-cell calcium signal. Magenta shading indicates when the cell was considered ‘on’. ARCOS groups ‘on’ cells in both space and time, and a circle fit to the ARCOS group indicates the approximate wave boundary. All waves fit in this work use the same code, outlined in Methods and available on GitHub.

**Figure SM3.**
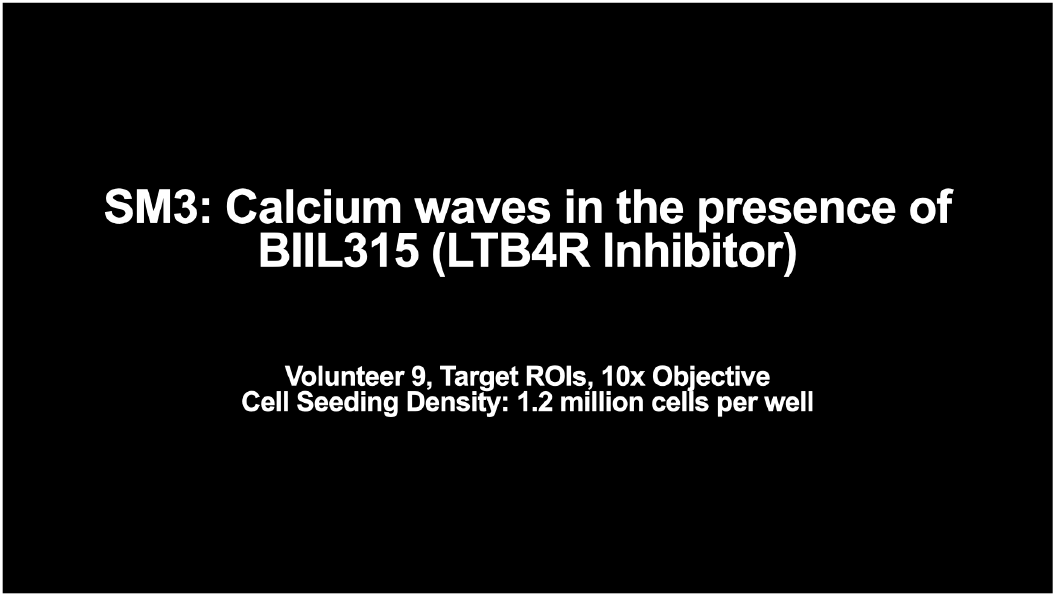
LTB4R signaling is required for rapid long-range swarming waves. Same imaging conditions to SM1 for control condition, target ROI displayed. Target well seeded with 1.2 million cells in both conditions. Both conditions shown use the same donor (Volunteer 9) and occur on the same day in adjacent wells. Imaging media is RPMI with 0.4% HSA and lacking phenol red. To block LTB4 sensing, neutrophils were resuspended in imaging media plus LTB4R inhibitor BIIL315 at a final concentration of 1 μM. Cells were incubated with or without inhibitor for 15min at room temperature and then seeded in a well for imaging. Calcium imaging is shown, then Hoechst imaging for both conditions.

**Figure SM4.**
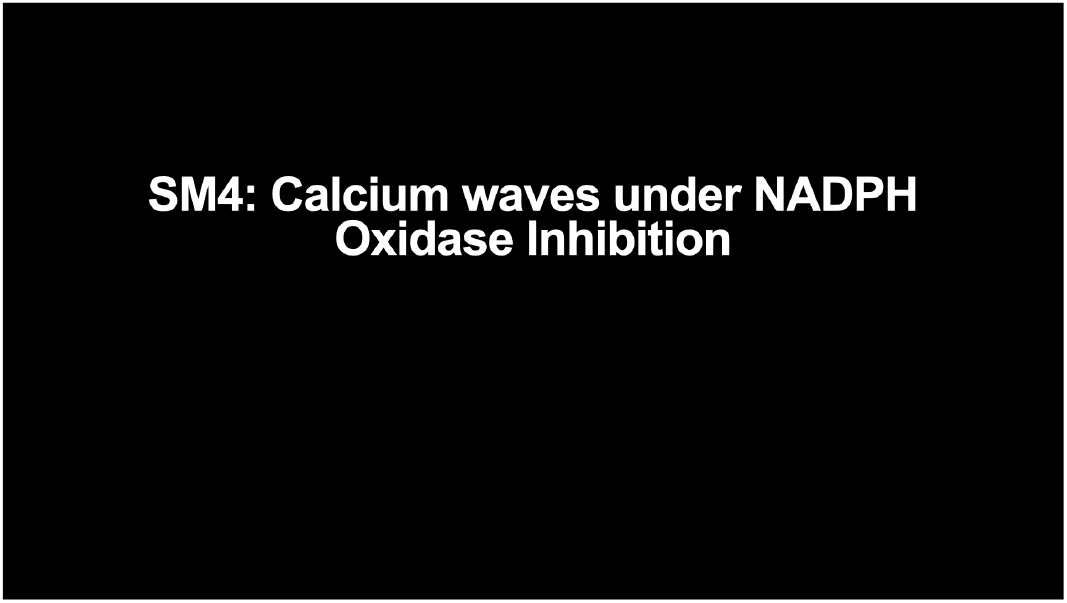
NADPH Oxidase activity is required for spatial restriction of swarming waves. First example is of healthy donor neutrophils treated with the NADPH Oxidase inhibitor DPI. Full frame ROI is displayed showing 4 targets. Neutrophils from Volunteer 19 were seeded at 1 million cells per well and imaged on the Ti2 Crest confocal system. To inhibit NADPH Oxidase activity, neutrophils were resuspended in imaging media plus DPI at a final concentration of 50 μM. Cells were pre-incubated with DPI for 15 min at room temperature and then seeded in a well for imaging. Calcium imaging is shown alongside Hoechst imaging. Next an ROI of this condition is played alongside CGD donor cells (which are deficient in NADPH Oxidase activity; Volunteer 338) swarming in the same assay. To minimize pre-activation in CGD neutrophils, cells were spun down to concentrate them and then left in R10 media and allowed to rest 15 min at room temperature before seeding into a well at a concentration of 1.2 million cells per well. Donor 338 is shown, and cells were imaged on the widefield microscope setup described in the Methods. Finally, the same CGD video is replayed with the calcium channel and Hoechst channel side-by-side.

**Figure SM5.**
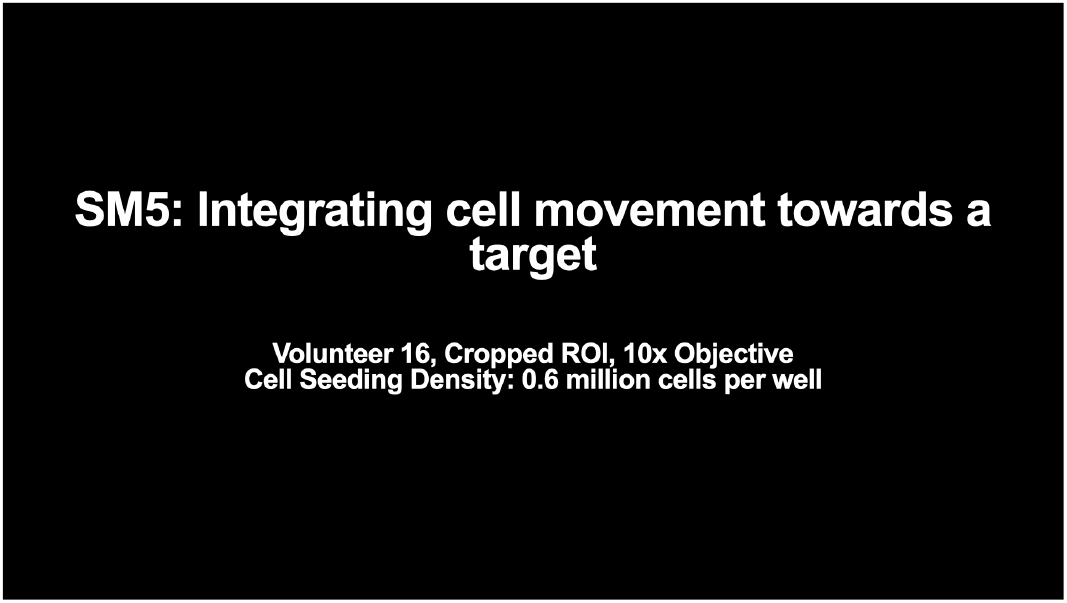
Multiple swarming waves elicit multiple rounds of neutrophil recruitment. Human neutrophils are seeded on to targets consisting of heat killed Candida albicans and recording begins. Calcium and Hoechst channels are imaged simultaneously similar to SM1. Cropped ROI, Volunteer 16, cells seeded at 600,000 cells per well. Three panels are shown. The left panel is the raw calcium imaging shown in greyscale. The middle panel is an output of both the wave tracking results superimposed on the cell tracking data. Movement that counts positively towards the integrated radial movement measurement is drawn in red, while movement that is subtractive to the sum is drawn in black. Movement that does not contribute more than *±* 1 *μ*m/5sec is drawn in grey, as this magnitude of movement is generated by noise in centroid calling. The right-most panel is a live graph that tracks the instantaneous radial velocity measurement synced with the movie and analysis panels.

**Figure SM6.**
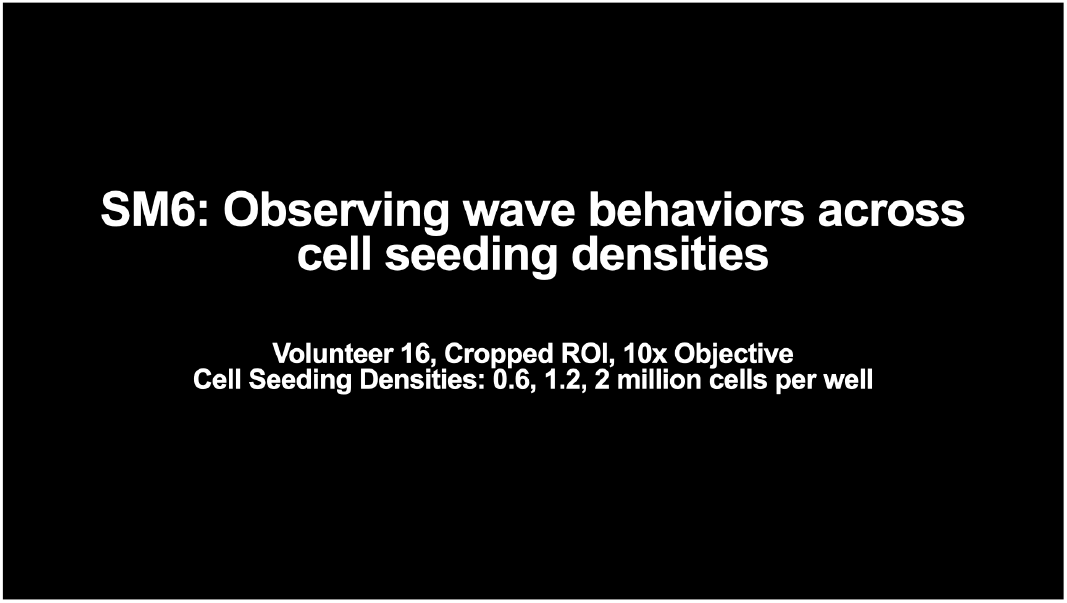
Neutrophils adjust swarming wave number/size to compensate for different initial cell densities. All examples are seeded onto targets of heat killed Candida albicans and recording begins. Cropped ROIs are shown. Cells all come from Volunteer 16 and were taken on the same day. Seeding densities are 0.6 million, 1.2 million, and 2 million cells per well. Calcium channels are shown with the wave detection overlayed. Next Hoechst channels of the same ROIs are shown with the wave detection overlayed.

**Figure SM7.**
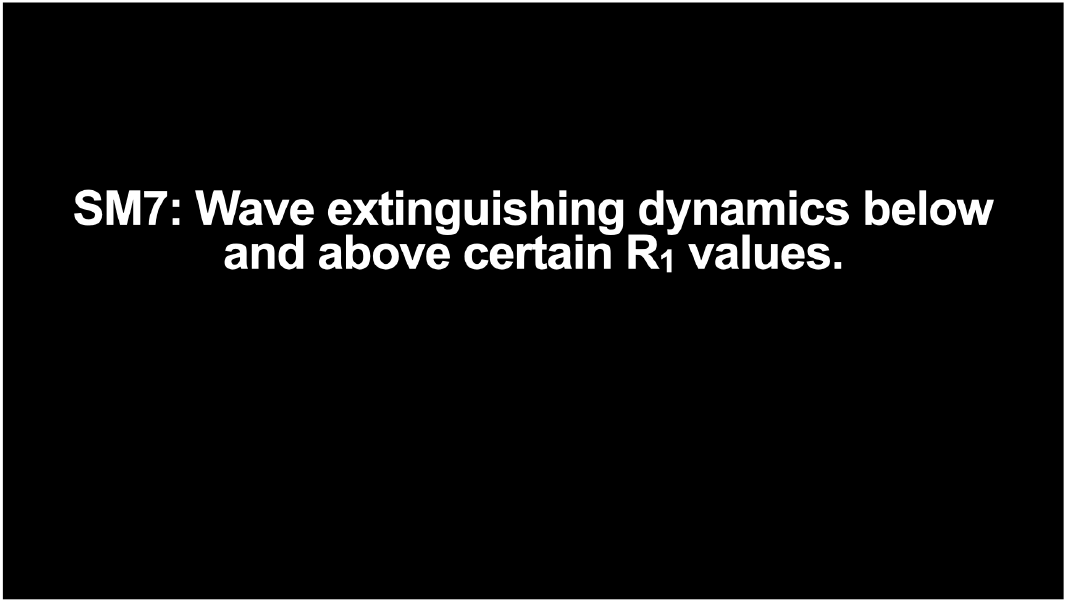
Wave extinguishing dynamics below and above various relay shuttoff *R*_1_ values and Hill coefficients. Some example simulations of Eq. 2 with different Hill coefficients. Higher Hill coefficients m,n,k=10,10,20 approaches the step function limit (Eq. 3) and results in a sharper wave stopping.

## Bibliography

Afonso, P. V., Janka-Junttila, M., Lee, Y. J., McCann, C. P., Oliver, C. M., Aamer, K. A., Losert, W., Cicerone, M. T., and Parent, C. A. LTB4 is a Signal-Relay molecule during neutrophil chemotaxis. Dev. Cell, 22(5):1079–1091, May 2012.

Alexander, N. J., Bozym, D. J., Farmer, J. R., Parris, P., Viens, A., Atallah, N., Hopke, A., Scherer, A., Dagher, Z., Barros, N., Knooihuizen, S. A. I., Saff, R. R., Pasternack, M. S., Thompson, R. W., Irimia, D., and Mansour, M. K. Neutrophil functional profiling and cytokine augmentation for patients with multiple recurrent infections: A case study. J. Allergy Clin. Immunol. Pract., 9 (2):986–988, Feb. 2021.

Aranyosi, A. J., Wong, E. A., and Irimia, D. A neutrophil treadmill to decouple spatial and temporal signals during chemotaxis. Lab Chip, 15(2):549–556, Jan. 2015.

Archambault, A.-S., Poirier, S., Lefebvre, J.-S., Robichaud, P.-P., Larose, M.-C., Turcotte, C., Martin, C., Provost, V., Boudreau, L. H., McDonald, P. P., Laviolette, M., Surette, M. E., and Flamand, N. 20-hydroxy- and 20-carboxy-leukotriene (LT) B4 downregulate LTB4-mediated responses of human neutrophils and eosinophils. J. Leukoc. Biol., 105(6):1131–1142, 2019.

Ataullakhanov, F. I., Guria, G. T., Sarbash, V. I., and Volkova, R. I. Spatiotemporal dynamics of clotting and pattern formation in human blood. Biochim. Biophys. Acta, 1425(3):453–468, Nov. 1998.

Barros, N., Alexander, N., Viens, A., Timmer, K., Atallah, N., Knooihuizen, S. A. I., Hopke, A., Scherer, A., Dagher, Z., Irimia, D., and Mansour, M. K. Cytokine augmentation reverses transplant recipient neutrophil dysfunction against the human fungal pathogen candida albicans. J. Infect. Dis., 224(5):894–902, Sept. 2021.

Birke, F. W., Meade, C. J., Anderskewitz, R., Speck, G. A., and Jennewein, H. M. In vitro and in vivo pharmacological characterization of BIIL 284, a novel and potent leukotriene b(4) receptor antagonist. J. Pharmacol. Exp. Ther., 297(1):458–466, Apr. 2001.

Chandra, V., Gal, A., and Kronauer, D. J. C. Colony expansions underlie the evolution of army ant mass raiding. Proc. Natl. Acad. Sci. U. S. A., 118(22), June 2021.

Chtanova, T., Schaeffer, M., Han, S.-J., van Dooren, G. G., Nollmann, M., Herzmark, P., Chan, S. W., Satija, H., Camfield, K., Aaron, H., Striepen, B., and Robey, E. A. Dynamics of neutrophil migration in lymph nodes during infection. Immunity, 29(3):487–496, Sept. 2008.

Dieterle, P. B., Min, J., Irimia, D., and Amir, A. Dynamics of diffusive cell signaling relays. Elife, 9: e61771, Dec. 2019.

Dinauer, M. C. Inflammatory consequences of inherited disorders affecting neutrophil function. Blood, 133(20):2130–2139, May 2019.

Donà, E., Barry, J. D., Valentin, G., Quirin, C., Khmelinskii, A., Kunze, A., Durdu, S., Newton, L. R., Fernandez-Minan, A., Huber, W., Knop, M., and Gilmour, D. Directional tissue migration through a self-generated chemokine gradient. Nature, 503(7475):285–289, Nov. 2013.

Fischer, J., Gresnigt, M. S., Werz, O., Hube, B., and Garscha, U. Candida albicans-induced leukotriene biosynthesis in neutrophils is restricted to the hyphal morphology. FASEB J., 35 (10):e21820, Oct. 2021.

Gagliardi, P. A., Grädel, B., Jacques, M.-A., Hinderling, L., Ender, P., Cohen, A. R., Kastberger, G., Pertz, O., and Dobrzynski, M. Automatic detection of spatio-temporal signalling patterns in cell collectives. July 2022.

Geiger, J., Wessels, D., and Soll, D. R. Human polymorphonuclear leukocytes respond to waves of chemoattractant, like dictyostelium. Cell Motil. Cytoskeleton, 56(1):27–44, Sept. 2003.

Gelens, L., Anderson, G. A., and Ferrell, J. E., Jr. Spatial trigger waves: positive feedback gets you a long way. Mol. Biol. Cell, 25(22):3486–3493, Nov. 2014.

Gordon, D. M. The ecology of collective behavior in ants. Annu. Rev. Entomol., 64:35–50, Jan. 2019.

Gregor, T., Fujimoto, K., Masaki, N., and Sawai, S. The onset of collective behavior in social amoebae. Science, 328(5981):1021–1025, May 2010.

Hamasaki, T., Sakano, T., Kobayashi, M., Sakura, N., Ueda, K., and Usui, T. Leukotriene B4 metabolism in neutrophils of patients with chronic granulomatous disease: phorbol myristate acetate decreases endogenous leukotriene B4 via NADPH oxidase-dependent mechanism. Eur. J. Clin. Invest., 19(4):404–411, Aug. 1989.

Henrickson, S. E., Jongco, A. M., Thomsen, K. F., Garabedian, E. K., and Thomsen, I. P. Noninfectious manifestations and complications of chronic granulomatous disease. J Pediatric Infect Dis Soc, 7(Suppl_1):S18–S24, May 2018.

Hopke, A., Scherer, A., Kreuzburg, S., Abers, M. S., Zerbe, C. S., Dinauer, M. C., Mansour, M. K., and Irimia, D. Neutrophil swarming delays the growth of clusters of pathogenic fungi. Nat. Commun., 11(1):2031, Dec. 2020.

Isles, H. M., Loynes, C. A., Alasmari, S., Kon, F. C., Henry, K. M., Kadochnikova, A., Hales, J., Muir, C. F., Keightley, M.-C., Kadirkamanathan, V., Hamilton, N., Lieschke, G. J., Renshaw, S. A., and Elks, P. M. Pioneer neutrophils release chromatin within in vivo swarms. Elife, 10: e68755, July 2021.

Jiwa, K., Ward, C., Parry, G., Brodlie, M., Lordan, J., Fisher, A., and Garnett, J. Immunohistochemical evidence of increased neutrophil swarms in cystic fibrosis lung removed at transplantation. Eur. Respir. J., 56(suppl 64):698, Sept. 2020.

Khazen, R., Corre, B., Garcia, Z., Lemaître, F., Bachellier-Bassi, S., d’Enfert, C., and Bousso, P. Spatiotemporal dynamics of calcium signals during neutrophil cluster formation. Proceedings of the National Academy of Sciences, 119(29):e2203855119, 2022.

Kienle, K., Glaser, K. M., Eickhoff, S., Mihlan, M., Knöpper, K., Reátegui, E., Epple, M. W., Gunzer, M., Baumeister, R., Tarrant, T. K., Germain, R. N., Irimia, D., Kastenmüller, W., and Lämmermann, T. Neutrophils self-limit swarming to contain bacterial growth in vivo. Science, 372(6548), June 2021.

Knooihuizen, S. A. I., Alexander, N. J., Hopke, A., Barros, N., Viens, A., Scherer, A., Atallah, N. J., Dagher, Z., Irimia, D., Chung, R. T., and Mansour, M. K. Loss of coordinated neutrophil responses to the human fungal pathogen, candida albicans, in patients with cirrhosis. Hepatol Commun, 5(3):502–515, Mar. 2021.

Kolaczkowska, E. and Kubes, P. Neutrophil recruitment and function in health and inflammation. Nat. Rev. Immunol., 13(3):159–175, Mar. 2013.

LaChance, J., Suh, K., Clausen, J., and Cohen, D. J. Learning the rules of collective cell migration using deep attention networks. PLoS Comput. Biol., 18(4):e1009293, Apr. 2022.

Lämmermann, T., Afonso, P. V., Angermann, B. R., Wang, J. M., Kastenmüller, W., Parent, C. A., and Germain, R. N. Neutrophil swarms require LTB4 and integrins at sites of cell death in vivo. Nature, 498(7454):371–375, June 2013.

Ley, K., Hoffman, H. M., Kubes, P., Cassatella, M. A., Zychlinsky, A., Hedrick, C. C., and Catz, S. D. Neutrophils: New insights and open questions. Sci Immunol, 3(30), Dec. 2018.

Mawhin, M.-A., Tilly, P., Zirka, G., Charles, A.-L., Slimani, F., Vonesch, J.-L., Michel, J.-B., Bäck, M., Norel, X., and Fabre, J.-E. Neutrophils recruited by leukotriene B4 induce features of plaque destabilization during endotoxaemia. Cardiovasc. Res., 114(12):1656–1666, Oct. 2018.

McDonald, P. P., McColl, S. R., Braquet, P., and Borgeat, P. Autocrine enhancement of leukotriene synthesis by endogenous leukotriene B4 and platelet-activating factor in human neutrophils. Br. J. Pharmacol., 111(3):852–860, Mar. 1994.

Molski, T. F., Naccache, P. H., Borgeat, P., and Sha’afi, R. I. Similarities in the mechanisms by which formyl-methionyl-leucyl-phenylalanine, arachidonic acid and leukotriene B4 increase calcium and sodium influxes in rabbit neutrophils. Biochem. Biophys. Res. Commun., 103(1): 227–232, Nov. 1981.

Muinonen-Martin, A. J., Susanto, O., Zhang, Q., Smethurst, E., Faller, W. J., Veltman, D. M., Kalna, G., Lindsay, C., Bennett, D. C., Sansom, O. J., Herd, R., Jones, R., Machesky, L. M., Wakelam, M. J. O., Knecht, D. A., and Insall, R. H. Melanoma cells break down LPA to establish local gradients that drive chemotactic dispersal. PLoS Biol., 12(10):e1001966, Oct. 2014.

Nauseef, W. M. Human neutrophils ≠ murine neutrophils: Does it matter? Immunol. Rev., 314 (1):442–456, Mar. 2023.

Ng, L. G., Qin, J. S., Roediger, B., Wang, Y., Jain, R., Cavanagh, L. L., Smith, A. L., Jones, C. A., Veer, M. d., Grimbaldeston, M. A., Meeusen, E. N., and Weninger, W. Visualizing the neutrophil response to sterile tissue injury in mouse dermis reveals a Three-Phase cascade of events. J. Invest. Dermatol., 131(10):2058–2068, Oct. 2011.

Park, S. A., Choe, Y. H., Park, E., and Hyun, Y.-M. Real-time dynamics of neutrophil clustering in response to phototoxicity-induced cell death and tissue damage in mouse ear dermis. Cell Adh. Migr., 12(5):424–431, Sept. 2018.

Parrish, J. K. and Edelstein-Keshet, L. Complexity, pattern, and evolutionary trade-offs in animal aggregation. Science, 284(5411):99–101, Apr. 1999.

Peiseler, M. and Kubes, P. More friend than foe: the emerging role of neutrophils in tissue repair. J. Clin. Invest., 129(7):2629–2639, June 2019.

Pettipher, E. R., Salter, E. D., Breslow, R., Raycroft, L., and Showell, H. J. Specific inhibition of leukotriene B4 (LTB4)-induced neutrophil emigration by 20-hydroxy LTB4: implications for the regulation of inflammatory responses. Br. J. Pharmacol., 110(1):423–427, Sept. 1993.

Poplimont, H., Georgantzoglou, A., Boulch, M., Walker, H. A., Coombs, C., Papaleonidopoulou, F., and Sarris, M. Neutrophil swarming in damaged tissue is orchestrated by connexins and cooperative calcium alarm signals. Curr. Biol., 30(14):2761–2776.e7, July 2020.

Reátegui, E., Jalali, F., Khankhel, A. H., Wong, E., Cho, H., Lee, J., Serhan, C. N., Dalli, J., Elliott, H., and Irimia, D. Microscale arrays for the profiling of start and stop signals coordinating human-neutrophil swarming. Nature Biomedical Engineering, 1(7):0094, July 2017.

Roxo-Junior, P. and Simão, H. M. L. Chronic granulomatous disease: why an inflammatory disease? Braz. J. Med. Biol. Res., 47(11):924–928, Nov. 2014.

Shaffer, B. M. Secretion of cyclic AMP induced by cyclic AMP in the cellular slime mould dictyostelium discoideum. Nature, 255:549–552, June 1975.

Siwicki, M. and Kubes, P. Neutrophils in host defense, healing, and hypersensitivity: Dynamic cells within a dynamic host. J. Allergy Clin. Immunol., 151(3):634–655, Mar. 2023.

Skoge, M., Yue, H., Erickstad, M., Bae, A., Levine, H., Groisman, A., Loomis, W. F., and Rappel, W.-J. Cellular memory in eukaryotic chemotaxis. Proc. Natl. Acad. Sci. U. S. A., 111(40): 14448–14453, Oct. 2014.

Song, Z., Huang, G., Chiquetto Paracatu, L., Grimes, D., Gu, J., Luke, C. J., Clemens, R. A., and Dinauer, M. C. NADPH oxidase controls pulmonary neutrophil infiltration in the response to fungal cell walls by limiting LTB4. Blood, 135(12):891–903, Mar. 2020.

Sun, D. and Shi, M. Neutrophil swarming toward cryptococcus neoformans is mediated by complement and leukotriene B4. Biochem. Biophys. Res. Commun., 477(4):945–951, Sept. 2016.

Sun, Y., Reid, B., Zhang, Y., Zhu, K., Ferreira, F., Estrada, A., Sun, Y., Draper, B. W., Yue, H., Copos, C., Lin, F., Bernadskaya, Y., Zhao, M., and Mogilner, A. Electric field-guided collective motility initiation of large epidermal cell groups. Mol. Biol. Cell, 34(5):ar48, May 2023.

Tweedy, L., Knecht, D. A., Mackay, G. M., and Insall, R. H. Self-Generated chemoattractant gradients: Attractant depletion extends the range and robustness of chemotaxis. PLoS Biol., 14(3):e1002404, Mar. 2016.

Uderhardt, S., Martins, A. J., Tsang, J. S., Lämmermann, T., and Germain, R. N. Resident macrophages cloak tissue microlesions to prevent Neutrophil-Driven inflammatory damage. Cell, 177(3):541–555.e17, Apr. 2019.

van Oss, C., Panfilov, A. V., Hogeweg, P., Siegert, F., and Weijer, C. J. Spatial pattern formation during aggregation of the slime mould dictyostelium discoideum. J. Theor. Biol., 181(3):203–213, Aug. 1996.

