## Supplemental Tables for "Self-extinguishing relay waves enable homeostatic control of human neutrophil swarming"

**Supplemental Table 1:** Healthy Control Donors drawn at UCSF under the IRB study number 21-35147 and used on the same day as blood draw. Full protocol in Supplemental Methods section. This cohort was used to generate all figures in this work **excluding** Figure 4F and Supplemental Figure 5B, D, F, and G.

| Donor #: | Sex: | Age: |
|---|---|---|
| **1** | M | 26 |
| **5** | M | 30 |
| **7** | M | 37 |
| **8** | M | 31 |
| **9** | F | 24 |
| **14** | M | 24 |
| **16** | F | 28 |
| **17** | F | 29 |
| **18** | M | 28 |
| **19** | M | 30 |

**Supplemental Table 2:** Healthy Control and CGD Donors drawn at the NIH campus in Bethesda, MD and shipped overnight to Boston, MA. Blood was processed immediately upon arrival and used that day. Full protocol in Supplemental Methods section. This cohort was used to generate Figure 4F and Supplemental Figure 5B, D, F, and G.

| Donor # | Sex: | Age: | Control/ CGD? | Protein Defect | Residual NADPH Oxidase Activity (nmoles/$10^7$ cells) |
|---|---|---|---|---|---|
| **337** | F | 48 | CGD | p47 | 3.05 |
| **338** | M | 22 | CGD | gp91 | 1.29 |
| **339** | M | 28 | Control | - | Not Tested |
| **340** | M | 27 | CGD | gp91 | 4.44 |
| **341** | M | 46 | Control | - | Not Tested |
| **348** | M | 41 | CGD | gp91 | 16.79 |
| **349** | M | 65 | Control | - | Not Tested |
