## Supplemental Theory Text for "Self-extinguishing relay waves enable homeostatic control of human neutrophil swarming"

**Self-Extinguishing Relay Model Supplemental Text**

To begin, we consider a reaction-diffusion model for active wave propagation. The cells should produce more diffusive activating ligand once they detect a sufficient amount of activating ligand. Without additional constraints on the positive feedback production of this ligand, it is challenging to construct a model that can propagate significantly longer than the diffusion limit and then stop. Therefore, we introduce a second molecule as an inhibitor to constrain the production of the activating ligand. Unlike the activator, the inhibitory molecule is assumed to be intracellular and non-diffusive, which is the known behavior of NADPH Oxidase with regards to LTB4 regulation in our own system (1).

In the continuous limit, we denote the activator molecule concentration as $L(r,t)$ and the inhibitor molecule concentration inside cells as $R(r,t)$. We can write the general form of the model as follows:

$$\begin{aligned} \frac{\partial L}{\partial t} &= D\nabla^2 L + f(L,R) \\ \frac{\partial R}{\partial t} &= g(L,R) \end{aligned} \tag{1}$$

Here, $f$ and $g$ can be arbitrary functions. However, to serve the activation and inhibition purposes, the activator production rate $f$ should depend positively on $L$ and negatively on $R$, while the inhibitor production rate $g$ should depend positively on $L$. We employ a Hill function form and consider the degradation of activator and inhibitor:

$$\begin{aligned} \frac{\partial L}{\partial t} &= D\nabla^2 L + a\rho \frac{R_1^m}{R^m + R_1^m} \frac{L^n}{L^n + L_1^n} - \gamma_L L \\ \frac{\partial R}{\partial t} &= b\frac{L^k}{L^k + L_2^k} - \gamma_R R \end{aligned} \tag{2}$$

In this model, $L_1, L_2$, and $R_1$ represent the dissociation constants for the activation of activator production, inhibitor production, and inhibition of activator production, respectively. The constant numbers $m, n$, and $k$ are Hill coefficients. Cell density $\rho$ and the production rate of the activator $a$ are presented together in this model, and we can define the product of them as the relay strength of signaling molecules. In the limit $R_1 \to \infty$, this model reverts to a single-molecule reaction-diffusion model with Hill function activation and simple decay, which has been extensively studied by prior works (2-7).

To further understand the wave-stopping mechanism, we can consider the limit $m, n, k \to \infty$ and $\gamma_L, \gamma_R \to 0$ to obtain a simplified model:

$$\begin{aligned} \frac{\partial L}{\partial t} &= D\nabla^2 L + a\rho\theta(R_1 - R)\theta(L - L_1) \\ \frac{\partial R}{\partial t} &= b\theta(L - L_2) \end{aligned} \tag{3}$$

In our experimental framework, a monolayer of cells (with a thickness denoted by $h$, approximately $10\mu m$) is positioned on a glass slide. The diffusion of molecules can extend up to the liquid layer's thickness ($H$, roughly 2000 $\mu m$). With a wave propagation speed, $v$, of about $2.5\ \mu m/s$ and the typical diffusion constant of LTB4, $D$, approximated at $125\ \mu m^2/s$, our set-up fulfills the conditions for the thick extracellular medium limit, represented by $H \gg \frac{D}{v} \gg h$ (6). By including the delta function of cell distribution over the $z$ axis, we derive the following equations:

$$\begin{aligned} \frac{\partial L}{\partial t} &= D\nabla^2 L + 2a\rho\theta(R_1 - R)\theta(L - L_1)\delta(z) \\ \frac{\partial R}{\partial t} &= b\theta(L - L_2) \end{aligned} \tag{4}$$

The extra factor of 2 originates from the semi-infinite environment for molecule diffusion since our simulations are done with infinite $z$ space. As prior research has demonstrated, in the absence of an inhibitor ($R_1 \to \infty$), this simplified model possesses several notable properties (6): 1. Under the traveling wave ansatz assumption, the wave speed can be analytically solved: $v = \frac{2a\rho}{\pi L_1}$, which does not depend on the diffusion constant $D$, as most diffusion in the 3D space will not trigger the relay process. 2. The concentration distribution at points distant from the wavefront of the threshold level can be approximated by: $L\left(\tilde{r} = r - vt \gg \frac{D}{v}\right) = \frac{a\rho}{v}\sqrt{\frac{D/v}{\pi\tilde{r}}}\exp(-v\tilde{r}/D)$, again under the traveling wave ansatz assumption. We can use these findings to better comprehend the wave-stopping mechanism when the inhibitor $R$ is present. As illustrated in **Supplemental Figure 3A**, which is a snapshot of the simulation results of the practical model, the concentration of the activator reaches $L_2$ and inhibitor starts to produce at position $r_1$ with a constant rate $b$. However, the inhibitor will not affect the production of activator until the inhibitor concentration reaches $R_1$, which is at position $r_2$ at the point in time which we are considering. If the activator wavefront $r_0$, at which the activator concentration is $L_1$, is behind $r_2$, the relay process will not be triggered due to inhibition effect, thus the activator cannot propagate further than a simple diffusion limit. Otherwise, if $r_2$ is behind $r_1$, it is possible that the activator in the range $(r_2, r_0)$ could trigger the production of more activators and thus lead to a persistent traveling wave.

Assuming a successful relay process, which in turn maintains a persistent traveling wave, the activation pulse duration $\tau$ at each position remains consistent: $\tau = (r_0 - r_2)/v$. Prior work found that the pulse duration needs to be longer than a minimum value to maintain the traveling wave (6):

$$\tau_m \approx 0.92D/v^2 = 0.92D(\pi L_1/2a\rho)^2 \tag{5}$$

under which the activator production on the 2D plane is not enough for compensating the diffusion in the 3D space. Intuitively, the distance $r_0 - r_2$ depends on the inhibitor production position $r_1$ and the travel distance before the inhibitor concentration is above the threshold to affect activator production $r_1 - r_2$. A lower $L_2$, a higher $L_1$, or a lower $R_1$ will effectively prohibit the relay process. (See **Supplemental Movie 7** for simulations with different parameter sets)

In the limit $\tau \gg D/v^2$, corresponding to a persistent relay, we can solve the positions $\tilde{r}_1, \tilde{r}_2$ approximately by assuming the traveling wave ansatz.

$$\begin{aligned} \tilde{r}_1 &= \frac{D}{v} f\left(\frac{L_1}{L_2}\right) \approx \frac{D}{v}\left(\ln\frac{L_1}{L_2} - \ln\frac{\pi}{2}\right), \text{when } \frac{L_1}{L_2} \gg 1 \\ \tilde{r}_2 &= \tilde{r}_1 - vR_1/b \\ v &= 2a\rho/\pi L_1 \end{aligned} \tag{6}$$

Which yields a dimensionless condition equivalent to $\tau = -\tilde{r}_2 \gg D/v^2$

$$f\left(\frac{L_1}{L_2}\right) \ll \beta = \frac{v^2 R_1}{bD} = \frac{4a^2\rho^2 R_1}{\pi^2 bDL_1^2} \tag{7}$$

As shown in **Supplemental Figure 3B**, our simulation verifies that the dimensionless $\beta$ is a plausible characteristic value for identifying wave-stopping behavior.

For comparison with the average result of multiple similarly sized waves, our model can produce similar wave-extinguishing behavior as observed in the experiments (**Figure 3D, Supplemental Figure 3c, d**).

**Citations**


1. Z. Song, G. Huang, L. Chiquetto Paracatu, D. Grimes, J. Gu, C. J. Luke, R. A. Clemens, M. C. Dinauer, NADPH oxidase controls pulmonary neutrophil infiltration in the response to fungal cell walls by limiting LTB4. *Blood*. 135, 891–903 (2020).

2. Keener, J. P. Propagation of Waves in an Excitable Medium with Discrete Release Sites. *Siam J Appl Math* 61, 317–334 (2000).

3. Kupferman, R., Mitra, P. P., Hohenberg, P. C. & Wang, S. S. Analytical calculation of intracellular calcium wave characteristics. *Biophys J* 72, 2430–2444 (1997).

4. Dawson, S. P., Keizer, J. & Pearson, J. E. Fire–diffuse–fire model of dynamics of intracellular calcium waves. *Proc National Acad Sci* 96, 6060–6063 (1999).

5. Mitkov, I., Kladko, K. & Pearson, J. E. Tunable Pinning of Burst Waves in Extended Systems with Discrete Sources. *Phys Rev Lett* 81, 5453–5456 (1998).

6. Dieterle, P. B., Min, J., Irimia, D. & Amir, A. Dynamics of diffusive cell signaling relays. *eLife* 9, e61771 (2020).

7. Dieterle, P. B. & Amir, A. Diffusive wave dynamics beyond the continuum limit. *Phys Rev E* 104, 014406 (2021).
