## Supplemental Methods for "Self-extinguishing relay waves enable homeostatic control of human neutrophil swarming"

**Neutrophil Isolation Protocol Draft:**

For all following uses, imaging media was first made with RPMI (w/o Phenol Red, 25mM HEPES, L-Glutamate) and 0.4% Human Serum Albumin (HSA)(Sigma, Cat: A5843). HSA was added directly to RPMI and then centrifuged at 500g for 5min until fully dissolved. The mix was then filtered with a .22um Steriflip filter (Millipore Sigma: SE1M179M6) before further use. Imaging media was always prepared fresh on the same day of imaging.

Blood specimens from patients were obtained with informed consent according to the institutional review board-approved study protocol at the University of California - San Francisco (Study #21-35147). Fresh samples of peripheral blood (2 tubes, 7mL each) from healthy adult volunteers were collected via a BD 23-gauge butterfly needle collection set (SKU: 23-021-022) into 10 ml BD Vacutainer EDTA tubes (SKU:366643). Volunteers were informed not to take ibuprofen, acetaminophen, or more than one drink of alcohol within 24hrs, class 1 or 2 antihistamines within 5 days, or aspirin within 7 days of blood draw. Blood was kept on a shaker at minimum setting and utilized within 2 hours of the draw. Neutrophils were isolated using the Stemcell EasySep Direct Human Neutrophil Isolation Kit (#19666) with the BigEasy magnet (#18001) per the manufacturer's protocol.

Isolated neutrophils were spun down at 200g for 5min and resuspended in a dye media consisting of imaging media plus 5ug/ml Hoechst 3334 (Invitrogen, Cat:H3570), and 1uM CalBryte 520 AM (AAT Bioquest, Cat: 20650). This cell suspension was incubated at room temperature in the dark for 15min, and then spun down at 200g for 5min. The dye medium was aspirated and replaced with an amount of cell culture media of RPMI (Gibco) with 10% heat inactivated FBS (Gibco) such that the final cell density was at or below 1x10^6 cells/mL. Purified neutrophils were then kept in polystyrene T25 flasks at 37C in a 5% CO2 environment until imaging. Cells were allowed to incubate for at least one hour before imaging began, and not used more than 5 hours after isolation. Allowing the cells to rest in culture before imaging helped ensure $Ca^{2+}$ signaling was more consistent and less noisy across volunteers.

CGD Cell Isolation:

Where CGD cells are used in this work the following changes to the above protocol were made. First, blood specimens were drawn from patients at the NIH campus for either control healthy donors or donors who are verified to have CGD. Samples were collected into 10ml BD Vacutainer EDTA tubes, and blood was shipped overnight in a temperature-controlled box using Phase 22 room temperature packs. Both CGD and healthy control blood were shipped together with each run to control for effects of shipping blood overnight. Blood was isolated in the same method as above as soon as it arrived in lab at around 10:30am each day experiments were performed. Cells were not used more than 5 hours after isolation, and all accumulation experiments all took place within 2 hours post isolation.

**Neutrophil Swarming Chip Protocol:**

Swarming arrays of C. albicans were prepared as described previously, with modifications (6, 7). Briefly, we used a microprinting platform (Picospotter PolyPico Galway, Ireland) to print a solution of 0.1% poly-l-lysine (Sigma-Aldrich) with ZETAG. For experiments, we printed arrays with either 1.0 mm or 1.5 mm spacing as indicated in an 8 well format on full sized No. 1.5H glass coverslips (Ibidi, Cat # 10812). Coverslips were dried and then left at room temperature until required. To attach an infection like material to these arrays, 8-well sticky-Slide attachments (Ibidi, Cat # 80828) were overlaid on the printed arrays. An overnight culture of live C. albicans yeast was heat killed at 90°C for 20 minutes before being washed and re-suspended in dH2O, then 750 µL of this suspension was added to each well and incubated for 5 minutes. Following incubation, wells were thoroughly washed out with dH2O to remove unbound targets from the glass surface. Wells were screened to ensure appropriate patterning of targets onto the spots with minimal non-specific binding before use. The well attachment was then removed, and the coverslips were stored at 4C until ready for use.

When preparing to use a patterned coverslip, the coverslip was first re-inspected for coating and target integrity. An Ibidi 8 well sticky-slide (Cat: 80828) was pressed firmly on to the slide, and a pipette tip was run along the bottom to ensure a proper seal was formed. The coverslip-well combo was then incubated in a 37C oven overnight. Next a 200ul mixture of imaging media plus 15 ug/ml Fibronectin from human plasma (Sigma, Cat: F0895) was pipetted into each well that was to be used that day. The slide was then incubated for 30min at 37C and washed 3x with 200ul/well of PBS (+/+ Ca/Mg). The final wash of PBS was left on the well until imaging.

**Neutrophil Swarming Imaging Protocol:**
Cells were imaged in one of two ways. Cells were imaged in the following way except when using CGD cells (See below). Neutrophils in culture were taken and placed into Fisherbrand LowRetention 1.5mL microcentrifuge tubes (Cat: 02681320) and spun down at 200g for 5min. Cells were resuspended at variable concentrations ranging from 3x10^6 - 10x10^6 cells/mL in a freshly made solution of imaging media. These cell solutions were then allowed to rest at room temperature for 15min. When the wait time had elapsed, the wash PBS was taken off the well to be imaged and 200ul of the neutrophil solution pipetted into the well. Imaging began as soon as imaging conditions could be verified after placing the cells into the well. All confocal microscopy data was collected using a Nikon Ti2-E body scope configured with a CrestOptics X-Light V2 confocal spinning disk system, a Lumencor Celesta light engine, Nikon 10x CFI Plan Apo Lambda objective, an Okobox temperature and C02 controlled environment, and a Photometrics Prime 95B sCMOS camera. All data was taken using the same levels of laser power from the 405nm and 488nm. The camera was run in a 2x2 binning mode with a set exposure time of 200ms for all channels. All movies are taken with a frame interval of 5 seconds between exposures unless otherwise indicated. All movies were taken at 37C with 5% CO2 for the duration of imaging.

Where indicated, this protocol was modified as follows to add the inhibitors used in this study. For the LTB4 inhibitor BIIL315 (Boehringer Ingelheim via opnMe), the drug first resuspended in DMSO at a stock concentration of 10mM. The stock was then diluted in resuspension media (RPMI+0.4%HSA) to a final concentration of 1uM before resuspending cells in the final step for imaging sample preparation. Cells were incubated in this drug-media solution for 15min before starting the experiment. The drug was kept in solution for the duration of imaging. For Diphenyleneiodonium chloride, or DPI (MedChemExpress, Cat: HY-100965), the drug was bought in a premixed solution of DMSO at a stock concentration of 10mM. The drug was diluted to a final concentration of 50uM in RPMI + 0.4%HSA before resuspending cells in the final step for imaging sample preparation. Cells were incubated in this drug-media solution for 15min before starting the experiment. The drug was kept in solution for the duration of imaging.

CGD Donor Cell Accumulation Imaging:
Neutrophils in culture were taken and placed into 1.5mL microcentrifuge tubes and spun down at 200g for 5min. Cells were resuspended at variable concentrations ranging from 3x10^6 - 10x10^6 cells/mL in a freshly made solution of imaging media. These cell solutions were then allowed to rest at room temperature for 15min. When the wait time had elapsed, the wash PBS was taken off the well to be imaged and 200ul of the neutrophil solution placed in the well. End point accumulation imaging was performed 60min after the wells were seeded with cells. Images were collected using a widefield florescence Nikon Ti-E body scope configured with a Nikon 10x Plan Flour objective equipped with a 37C temperature and 5% CO2 environment chamber.

CGD Donor Calcium Imaging:
Neutrophils in culture were taken and placed into 1.5mL microcentrifuge tubes and spun down at 200g for 5min. To avoid extra potential activation, cells were resuspended in the same media they were cultured in, RPMI + 10% FBS. These cells were left to rest for 15min at room temperature, and then added to the well and imaged immediately. Timelapse imaging was performed on a Nikon Ti-E body widefield florescence scope, with a 10x Nikon objective, and wells were inside a stage top 37C incubation unit.

**Image Analysis:**
All image analysis presented in this work was achieved via the use of both Fiji (ImageJ) and Python. Images were pre-processed when needed to split one field of view into 4 quadrant ROIs to simplify the downstream Python analysis. All python analysis for each graph is provided as open-source code in the form of both scripts and Jupyter notebooks on Github (https://github.com/strickland-ev/swarming-self-extinguishing-relay-publication). A brief description of each data analysis tool used is provided in plain text below. The raw data for all graphs generated will be provided as a zip file on Box or the publisher website. We also intend to publish our raw data, code, and all files that went into making these figures and movies on Zenodo or comparable platform at the time of publication.

Movie Flat Field Correction:

While data used for image analysis pipelines was not normally flat field corrected due to the local nature of the measurements taken, images shown in some of the movies were flat field corrected with the following simple protocol using FIJI. Movies shown in SM1, 3, and 4 were flat field corrected by taking the entire frame and blurring each timepoint with a gaussian blur filter set to 150px radius. A time stack projection was then made where the median value was taken across all timepoints for each pixel in XY. This median projection of the blurred stack was taken as the background and subtracted from the original image stack. This process was performed independently on each florescence channel. The macro used to automate some of this process will be included in the data posted alongside this publication.

ARCOS Wave Tracking:
See code posted on Github for a more direct explanation of the methods used and the code used for each figure. Broadly, to track cells trackpy was used to link frames where nuclei were identified via a custom StarDist model. The ARCOS algorithm was used to group like calcium events. Further analysis of cell tracks was done via custom python code included in the Github jupyter notebooks.
